# Supervised learning of protein variant effects across large-scale mutagenesis datasets

**DOI:** 10.1101/2025.04.02.646878

**Authors:** Thea K. Schulze, Lasse M. Blaabjerg, Matteo Cagiada, Kresten Lindorff-Larsen

## Abstract

The increasing availability of data from multiplexed assays of variant effects (MAVEs) enables supervised model training against large quantities of experimental data to learn sequence-function relationships. Variant effect scores from MAVEs can, however, be influenced by the experimental method to create experiment-to-experiment differences in the mapping from molecular level-variant effects to MAVE readout, which presents a challenge for supervised learning across datasets. We here propose a framework for performing supervised learning with MAVE data that takes the influence of the experimental protocol into account, thus enabling variant effects to be learned across datasets produced in independent experiments. We apply the framework to train a model against variant effect scores collected with VAMP-seq, a MAVE technique that quantifies the steady-state cellular abundance of protein variants. We show that mapping variant abundance to VAMP-seq readout in a dataset-specific manner during model training improves the learned abundance model and moreover allows the learned model to predict variant effects on an interpretable scale. Our work highlights the importance of validating MAVE results with low-throughput methods to facilitate MAVE score interpretation and supervised model training.

## Introduction

Large-scale mutagenesis experiments, such as multiplexed assays of variant effects (MAVEs), make it possible to quantitatively characterise the effects of thousands of amino acid residue substitutions on protein stability and function in a single experiment. Often, all possible single residue substitutions in a protein are assayed in such experiments, thereby facilitating thorough analysis of the importance of different residue positions and types for maintaining the structural and functional integrity of a protein. Additionally, as many proteins have now been studied with MAVE methods (Esposito et al., 2019; Rubin et al., 2021; Beltran et al., 2025), several large-scale mutagenesis datasets can be combined to (*i*) analyse effects of residue substitutions across protein backgrounds (Gray et al., 2017; Dunham and Beltrao, 2021; Schulze and Lindorff-Larsen, 2024; Beltran et al., 2025), (*ii*) benchmark the ability of existing computational tools to classify and predict variant effects (Livesey and Marsh, 2023; Gerasimavicius et al., 2023; Notin et al., 2023) and (*iii*) train models that aim at predicting variant effects in proteins that were not seen during training (Gray et al., 2018; Høie et al., 2022; Jagota et al., 2023; Fu et al., 2023). Development of such predictors is, among other things, motivated by the fact that experimental mutagenesis data currently far from cover all possible residue sites and substitution types in the human proteome (Tabet et al., 2022; Fowler et al., 2023), which limits clinical and mechanistic interpretation of observed variants (Fayer et al., 2021) and thus potentially disease treatment or drug development (Bernier et al., 2004; Mighell and Lehner, 2024). Moreover, accurate predictors have potential applications in protein engineering (Freschlin et al., 2022).

The use of MAVE data for training predictive models has already been explored in a number of studies, revealing that variant effects can indeed be learned from these data (Gray et al., 2018; Gelman et al., 2021; Munro and Singh, 2021; Høie et al., 2022). This is not only the case when models are trained from scratch; fine-tuning of large, pre-trained architectures and transfer learning with embeddings from such architectures against MAVE data have resulted in variant effect estimators that offer improved performance over zero-shot predictions from the original models (Hsu et al., 2022a; Jagota et al., 2023; Dieckhaus et al., 2024; Lafita et al., 2024). Variant effect predictions from models trained on MAVE data are, however, often non-perfect, and extrapolation to effects on proteins outside the training set can be challenging. There could be several reasons for this; here, we focus on reasons related to how MAVE data are used for training and validation of the predictive models.

A variety of experimental approaches can be classified as MAVEs. These approaches have in common that the impact of many different sequence variants on a molecular phenotype of interest are measured in parallel using a combination of high-throughput sequencing and designed, often cell-based, expression systems that link genotype and the corresponding molecular phenotype to a cellular phenotype that can easily be read out by DNA sequencing (Fowler and Fields, 2014; Weile and Roth, 2018; Geck et al., 2022). MAVEs have been used to study the effects of variants on many different molecular phenotypes, including catalytic eiciency, ligand binding ainity and cellular stability. The experimental readout, which might for example be cell growth or fluorescence intensity, can change from assay to assay, even when the molecular phenotype is kept constant (Faure et al., 2022; Matreyek et al., 2018). As a result, MAVE data are highly diverse, even when variation in experimental conditions is not considered.

In a few studies, variant effects have been measured for many (typically small) protein domains in single experiments (Rocklin et al., 2017; Tsuboyama et al., 2023; Beltran et al., 2025). However, most MAVEs are performed for one protein at a time, and supervised training of models thus often rely on combining multiple independent datasets reporting on variant effects in different proteins (Gray et al., 2018; Høie et al., 2022; Jagota et al., 2023; Fu et al., 2023). The combination of several MAVE datasets to facilitate learning across proteins often results in a relatively heterogeneous collection of variant effect scores and might for example mix scores reporting on different molecular phenotypes in different organisms under different experimental conditions. Moreover, rescaling and normalisation of individual datasets is typically needed to bring scores from different experiments onto a comparable scale. It is, however, not clear that such approaches necessarily make scores from different assays compatible, as one could for example imagine that the mapping from molecular phenotype to MAVE readout depends on the readout type. Variant effect score combination might thus add noise to supervised training of new predictors for several reasons.

Here, we set out to develop a variant effect model specifically for predicting effects of residue substitutions on cellular abundance. By focusing on a single molecular phenotype, we limited the possible heterogeneity in a combined variant effect score training dataset. The cellular concentration or abundance of a protein is regulated by proteostasic processes (Labbadia and Morimoto, 2015), but can be affected by single residue substitutions, and a significant amount of pathogenic mutations in proteins cause disease because they reduce cellular abundance (Yue et al., 2005; Sahni et al., 2015; Jänes et al., 2024; Cagiada et al., 2025; Beltran et al., 2025). The steady-state abundance of a protein in a cell is a balance between protein synthesis and degradation rates; here, we focus on whether substitutions affect protein degradation. Changes in cellular abundance due to changed degradation behaviour correlate with measurements and predictions of folding free energy changes for variants of many proteins (Nielsen et al., 2017; Abildgaard et al., 2019; Matreyek et al., 2018; Suiter et al., 2020; Gerasimavicius et al., 2023; Grønbæk-Thygesen et al., 2024; Clausen et al., 2024). Mechanistically, structural destabilisation that causes global or local unfold-ing of the protein structure is thought to expose (quality control) degrons to the protein quality control system or degradation machinery of the cell and thereby reduce abundance through increased degradation rates (Pey et al., 2007; Stein et al., 2019; Kampmeyer et al., 2022). Residue substitutions might also result in changed protein levels for other reasons (Correa Marrero and Barrio-Hernandez, 2021), for example by interfering with other degron types, post-translational modification patterns (Vazquez et al., 2000) or the ability to form protein complexes (Suiter et al., 2020; Grønbæk-Thygesen et al., 2024; Jänes et al., 2024).

To learn a general model for how substitutions affect cellular abundance, we trained a predictor against six independent MAVE datasets. We restricted our training data to consist only of variant effect scores collected with variant abundance by massively parallel sequencing (VAMP-seq), a MAVE method that estimates the impact of substitutions on the steady-state cellular abundance of a protein (Matreyek et al., 2018). In a VAMP-seq experiment, protein variants are expressed fused to a fluorescent protein tag in cultured human cells. Each cell expresses a single variant, and cellular fluorescence intensity thus reports on the concentration or abundance of the expressed variant. Variant-expressing cells are sorted with fluorescence-activated cell sorting (FACS) into a discrete number of populations or bins covering different fluorescence intensity intervals. By sequencing the sorted populations, the number of times each variant occurs in each population can be counted, and these counts can be transformed into a set of scores that indicates how every assayed substitution affects abundance (Matreyek et al., 2018). A typical approach in previous VAMP-seq work has been to FACS-sort cells into four discrete populations so that each population would end up containing 25% of the total number of cells in the experiment (Matreyek et al., 2018, 2021; Amorosi et al., 2021; Suiter et al., 2020; Grønbæk-Thygesen et al., 2024; Clausen et al., 2024). The strategy of sorting and sequencing is useful as it facilitates parallel assessment of thousands of variant effects. However, the choice to sort cells into equally populated bins has important implications for how the resulting data should be interpreted; when cells are sorted this way, the final variant effect scores obtained will depend on the composition of the library of cells being sorted, and the experiment thus ultimately produces relative scores that inform on the ranking of variant effects rather than on absolute effects. Consider, for example, a scenario in which two cell libraries of equal size express the same variants of the same protein. The only difference between the libraries is that one of them includes a large number of cells that express the wild-type protein and the other one does not. When the cells from the two libraries are sorted into four equally large populations, some of the cells that express non-synonymous variants will end up in different sorted fractions, because one sorting must accommodate the wild-type protein-expressing cells. As a result, the variant effect score measured in one experiment will not match the variant effect score measured in the other experiment for some variants. It is thus not necessarily trivial to combine variant effect scores from two different VAMP-sequences experiments.

To perform supervised learning against six independent VAMP-seq datasets at once, we therefore took the following approach. We first analysed the datasets and found that there is indeed experiment-to-experiment variation in how the variant effect scores produced in high throughput relate to low-throughput measurements of cellular protein levels for selected variants. This presents a challenge for learning across the datasets. Then, to overcome this challenge, we developed a learning framework that models the experimental process and thereby allows dataset- or experiment-specific properties to be taken into account during supervised model training. We applied the framework to the six VAMP-seq datasets and found that our modelling of the experimental process makes it possible to learn additional patterns in the training data, but also introduces new modelling challenges. An advantage of our framework is that it facilitated development of an abundance predictor that outputs variant effects on an interpretable scale, even though it was trained against VAMP-seq data. Our work generally shows that even when the molecular phenotype and MAVE type are kept constant to limit variant effect score heterogeneity in training and validation data, supervised learning across MAVE datasets needs to be done carefully with attention towards the experimental data that is being used. With appropriate modifications, the supervised learning framework developed here might be useful to learn variant effects across other types of MAVE data.

## Results and Discussion

### Experiment-specific mappings from low- to high-throughput abundance scores

The VAMP-seq method has been used to measure the impact of single residue substitutions on cellular abundance for several proteins, including both soluble proteins and membrane proteins. We have recently combined and analysed VAMP-seq data for six soluble proteins (Schulze and Lindorff-Larsen, 2024), namely PTEN (Matreyek et al., 2018, 2021), TPMT (Matreyek et al., 2018), CYP2C9 (Amorosi et al., 2021), NUDT15 (Suiter et al., 2020), ASPA (Grønbæk-Thygesen et al., 2024) and PRKN (Clausen et al., 2024), and we here set out to use this combined VAMP-seq dataset for supervised learning. Aggregation of data from the six VAMP-seq experiments gives a combined dataset containing ca. 32,000 single residue substitution variant effect scores.

Importantly, the cellular abundance of variants of the six proteins in the combined dataset was measured in individual VAMP-seq experiments, one experiment per protein, and very similar meth-ods were used to collect the individual datasets. All six experiments quantified variant effects using a library of cells in which each cell expressed a single protein variant labelled with green fluorescent protein (GFP). GFP-tagged variants and mCherry were co-expressed via an internal ribosome entry site, thus allowing for normalisation to the expression levels of individual cells. The library of variant-expressing cells was in each experiment sorted into four equally populated bins based on GFP:mCherry fluorescence intensity ratios using FACS. Then, a variant effect score was obtained for every variant by calculating a weighted average of the variant frequency across the four bins after the bins were sequenced. These scores were finally normalised so that scores of 0 and 1 would respectively indicate nonsense variant-like and wild-type variant-like cellular protein levels. We will refer to these variant effect scores as VAMP-seq or high-throughput scores below. We note that the VAMP-seq method has produced variant effect scores with relatively high reproducibility across replicates for all six datasets considered here.

When the six VAMP-seq experiments were performed, GFP:mCherry fluorescence intensity ratios from populations of cells that all expressed the same protein variant were measured for a few (between 11 and 24) variants in low throughput, that is, for one variant at a time, to thereby estimate the cellular protein levels of the selected variants in a more direct fashion and hence validate the results of the high-throughput VAMP-seq experiment. In the ideal case, the low- and high-throughput variant effect scores would be highly linearly correlated. However, as described above, the absolute scores obtained from a VAMP-seq experiment are affected by, among other things, the composition of the sorted library of cells. Consequently, there is no guarantee that the resulting data will necessarily follow this ideal trend or even follow similar trends across independent experiments.

To evaluate and compare the extent to which VAMP-seq scores from the six datasets correlate with low-throughput measurements of cellular protein levels, we collected the previously measured low-throughput data and plotted the low- and high-throughput scores against each other (Fig. 1A). Spearman’s rank correlation coeicient, *r_s_*, is relatively high between the two score types for several proteins, thus indicating that the VAMP-seq scores provide relatively accurate estimates of the ranking of variant effects for those proteins (Fig. 1B). The linear correlation, *r*, between the low- and high-throughput scores is, however, lower than the rank correlation for most datasets (Fig. 1B). Importantly, the way in which the data deviate from linearity depends on the dataset, and there is considerable experiment-to-experiment variation in the way that the low- and high-throughput scores relate to each other.

**Figure 1.**
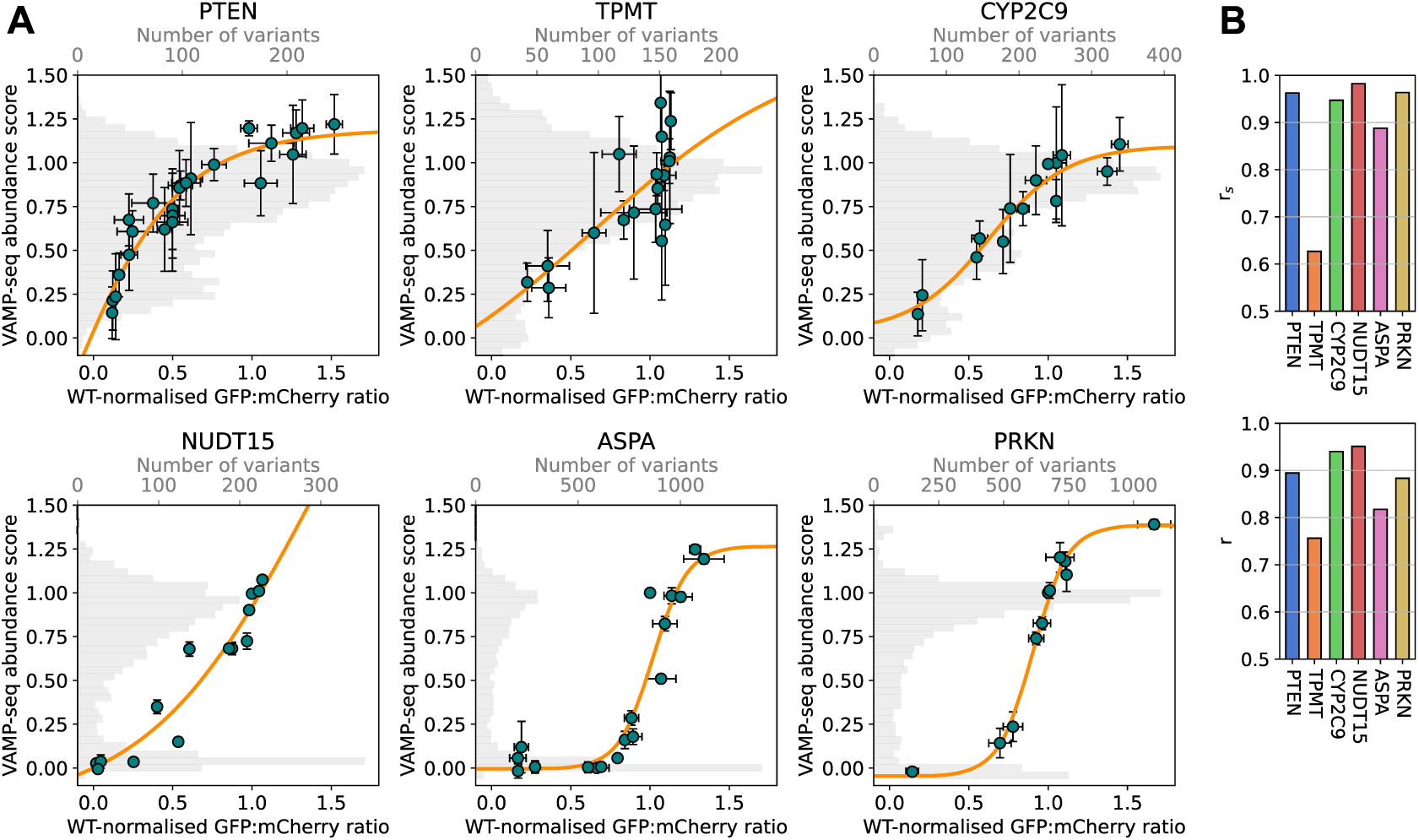
Experiment-specific mappings from low- to high-throughput abundance scores. (A) Cellular abundance was measured for thousands of variants in high throughput with the VAMP-seq method, and these data were validated for a few selected variants with low-throughput measurements of cellular protein levels in the form of fluorescence intensities from populations of cells that all expressed a single variant. The green data points show how the low-throughput abundance scores (WT-normalised GFP:mCherry ratios) relate to the high-throughput abundance scores (VAMP-seq abundance scores) for the validation variants for each of the six proteins in our combined dataset. The error bars on the WT-normalised GFP:mCherry ratios indicate the standard error of the mean, whereas the VAMP-seq score error bars are standard deviations. Error bars are missing on NUDT15 low-throughput scores. We fit small multilayer perceptrons (MLPs) to the validation data points to obtain standard curves (shown in orange) that for each protein relate the high-throughput VAMP-seq scores to the low-throughput measurements. The distributions of all available VAMP-seq scores for single residue substitution variants of the six proteins are shown in the plot backgrounds in grey with the corresponding x axes on top of the plots. The shapes of the distributions can be explained by the shapes of the standard curves. For example, many ASPA variants have very low abundance scores because variants with a wide range of low-throughput scores were sorted into the same fraction in the VAMP-seq experiment (***Grønbæk-Thygesen et al., 2024***). (B) Spearman’s rank correlation coeicient, *r_s_*, and Pearson’s correlation coefficient, *r*, between low- and high-throughput abundance scores for the six individual datasets.

This variation for example becomes clear when comparing the ASPA, PRKN and PTEN data. ASPA and PRKN variants that cover a wide range of cellular protein levels are assigned variant effect scores close to 0 in the high-throughput experiments, while WT-like variants of the two proteins spread out across the high-throughput score distributions. Conversely, several PTEN variants with varying intracellular concentrations have WT-like scores in the high-throughput experiment (Fig. 1A). Similar VAMP-seq scores from two separate experiments might thus translate to very different protein levels as measured in low throughput, meaning that the VAMP-seq scores do not necessarily report on variant cellular abundance in a consistent manner across experiments. While this is less of a problem when analysing individual proteins, this implies that VAMP-seq scores cannot easily be combined to perform supervised learning across the datasets unless the experiment-specific relations between low- and high-throughput scores are taken into account. We propose that this can be done by using standard curves that relate the two score types to each other and that such standard curves can be fitted using the available validation data (Fig. 1A). Alternatively, standard curves might be obtained from more elaborate modelling of the FACS process that takes the composition of the sorted cell library into account (Peterman and Levine, 2016).

### Model architecture for supervised learning across VAMP-seq datasets

We developed a framework for learning variant effects on cellular abundance across VAMP-seq datasets and hence proteins that integrates the variant effect prediction task with modelling of the experimental process in a modular manner (Fig. 2). The framework is built on the hypothesis that the WT-normalised GFP:mCherry ratios measured in low throughput represent the true variant impact on abundance more accurately than the count-based VAMP-seq scores do. We thus ultimately aimed at developing a model that would be able to predict variant abundance as measured in low-throughput experiments. However, since low-throughput measurements were available for only a small number of variants and would, as a result, likely not be suicient for model training, we constructed the framework so that the thousands of VAMP-seq scores measured in high throughput could instead be used as a means to learning the low-throughput target.

**Figure 2.**
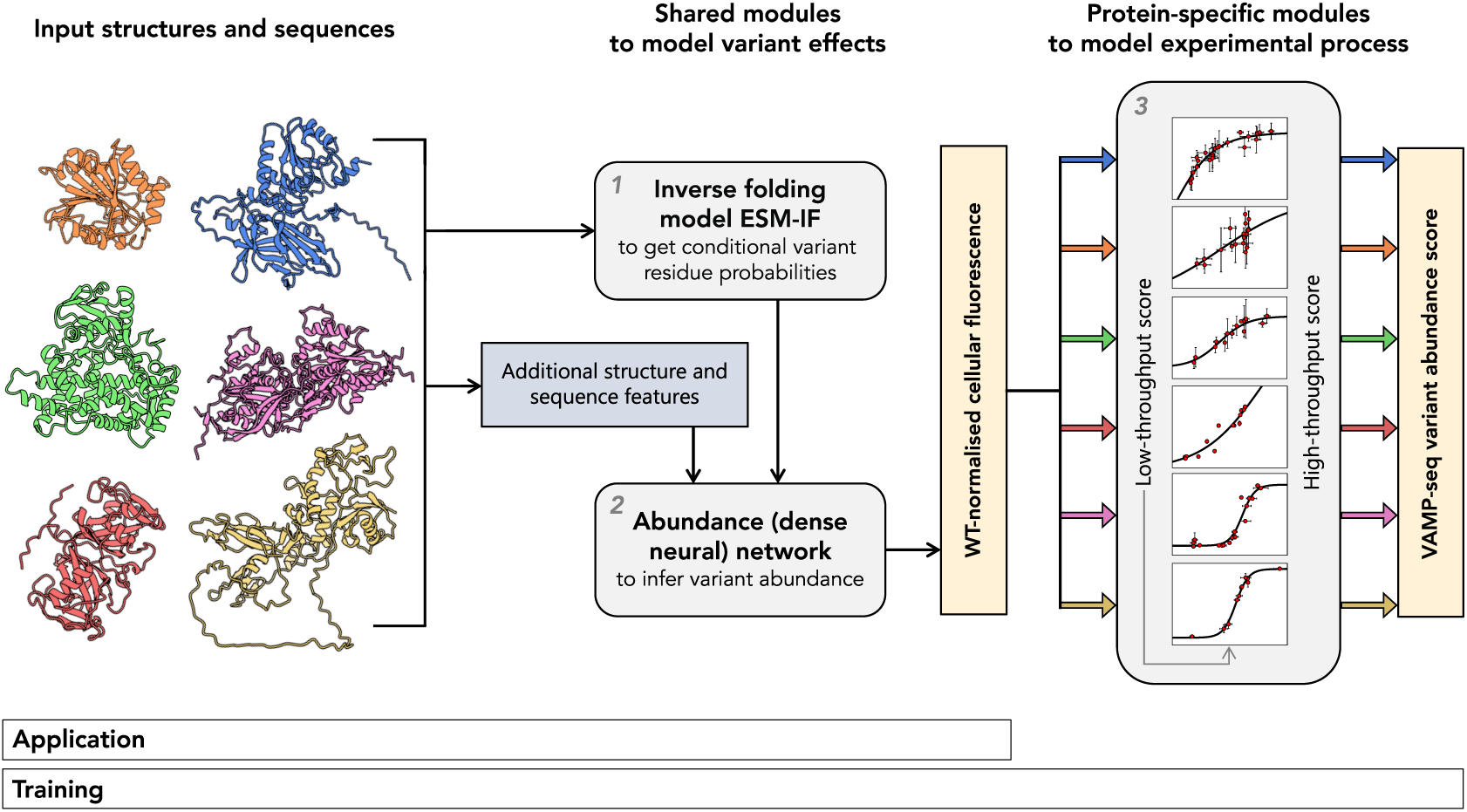
Framework for supervised learning across VAMP-seq datasets. The learning framework consists of three neural network modules (shown in grey boxes and with numbers *1*, *2* and *3*) with different information processing tasks. The first neural network module (*1*) is ESM-IF (***Hsu et al., 2022b***), which, given a protein sequence and structure, can be used to calculate probabilities for all possible single residue substitutions at the target residue site. The second neural network (*2*) is the actual abundance model that processes the output from ESM-IF along with other sequence- and structure-derived features to predict variant abundance in the form of WT-normalised (GFP:mCherry) fluorescence intensity ratios as measured in a low-throughput experiment. If the abundance network has already been trained, the output from this network would constitute abundance score predictions for a new protein. To train the abundance network using VAMP-seq data from multiple individual experiments, predicted WT-normalised fluorescence intensity scores are passed through the framework’s third and final module (*3*), which consists of the experiment- or protein-specific standard curves shown in orange in Fig. 1. Transformation of predicted low-throughput scores through the standard curves gives output in the form of VAMP-seq scores. A loss between predicted and experimental VAMP-seq scores can be calculated and used to update the parameters of the abundance network (*2*). Information runs through the framework for a single variant of a single protein at a time, although a full structure is needed to generate the input features to the second module. The protein structures to the left are TPMT (orange), PTEN (blue), CYP2C9 (green), ASPA (pink), NUDT15 (red) and PRKN (yellow).

In our framework, the target during training is thus the VAMP-seq score of a single residue substitution variant of an input protein. The VAMP-seq score is predicted from the input protein’s sequence and structure from a series of neural network modules that are used to (*1*) parse the input structure, (*2*) integrate a number of structure and sequence features to estimate the effect of a substitution on cellular abundance as it would be measured in a low-throughput experiment, and (*3*) model the experimental process by transforming the estimated low-throughput score to a VAMP-seq score via the experimentally informed relation between the two score types. This latter transformation is protein- or experiment-specific and accounts for experiment-to-experiment variation in low- to high-throughput score mappings during training. Thus, with this transformation, we should be able to train the second module of the framework, the low-throughput score predictor, using the high-throughout data in spite of the differences in high-throughout score meanings across datasets.

We now elaborate on the details of each of the modules that make up our learning framework. In the first part of the model (Fig. 2.1), features are extracted from the protein sequence and structure, primarily by using ESM-IF, the ‘inverse folding’ model from the Evolutionary Scale Model family (Hsu et al., 2022b). ESM-IF is a generative model developed to suggest amino acid sequences that would fit a given protein backbone template and was trained in a self-supervised fashion to recover native sequences of natural proteins. Given both the structure and sequence of a protein, the model can be used to estimate the likelihood of each of the 20 standard amino acids to occur at any given position in the protein. For all variants of the six proteins, we used ESM-IF to estimate the likelihood of the variant sequence given the wild-type structure and found that the ratios between the variant and wild-type sequence likelihoods correlate well with the VAMP-seq scores (Fig. S1). Based on that, we used the ESM-IF likelihood ratios (which we refer to as ESM-IF scores below) as features for downstream modelling.

In our framework, ESM-IF scores for the wild type residue and the 19 possible substitutions at the target variant residue position are fed into a downstream dense neural network, the abundance network (Fig. 2.2), together with one-hot encoded representations of the wild-type and variant residue types and a number of features reporting on the local and (sub)global structure and sequence environment of the target variant (Fig. S2). The additional features were engineered to provide information about the thermodynamic and cellular stability of the target variant and are discussed further below. The task of the abundance network is to integrate the input features to predict the impact of the variant on cellular abundance in the low-throughput space, that is, to predict variant impact on WT-normalised cellular fluorescence levels.

To be able to learn the parameters of the abundance network using the high-throughput data, the output of the network is passed through a protein- or experiment-specific module that transforms the low-throughput score prediction into a high-throughput VAMP-seq score using experimentally-informed standard curves (Fig. 2.3). We obtained the standard curves by fitting small networks to describe the mappings from low- to high-throughput scores (orange curves in Fig. 1A), and our modelling thus depends on having both low- and high-throughput scores available for all six systems. This third module of the framework is, however, only needed for training purposes and should thus be removed when abundance predictions are made for new proteins.

### Improved abundance score predictions from standard-curve-based models

We trained and evaluated the learning framework using leave-one-protein-out cross-validation and quantified the performance of the framework by calculating the mean absolute error between the experimental and predicted VAMP-seq scores of the protein that was left out for validation.

We also calculated *r_s_* between the experimental VAMP-seq scores and the abundance network output to evaluate trained models in a way that does not depend on the standard curves. We generally assumed that if a model could predict abundance scores measured in high throughput more accurately than another model, the accuracy of the former model should also be highest in the low-throughput score space. Although we aimed to train a low-throughput-score predictor, we first evaluated the utility of including standard curves as part of the learning framework for high-throughput-score prediction.

Specifically, to study the impact on the learning process of including the standard curves in the framework, we trained two sets of leave-one-protein-out models. First, we trained a set of leave-one-protein-out models in which the entire learning framework was used, that is, where the standard curves were used to transform output of the dense network before calculating the loss between predictions and experimental data. We then also trained a set of models in which the dense network was directly tasked with predicting the high-throughput scores, that is, in which we omitted the standard curves from the framework (Fig. S3). During the training of both sets of models, only the parameters of the abundance network were allowed to be modified; when the standard curve models were present in the framework downstream of the abundance network, their parameters were frozen, and ESM-IF parameters were also never adjusted.

After optimising hyperparameters for both sets of models, we observed consistent improvements in the ability to predict the high-throughput abundance scores when the standard curves were included during training and validation (Fig. S4, S5). Only when PTEN was used as validation protein did inclusion of the standard curves seem to lower the model accuracy, with the decrease in accuracy being quite significant (Fig. S4, S5). Motivated by this observation (see also discussion of PTEN data in the supplementary material), we retrained both sets of models leaving PTEN VAMP-seq scores out of the training data (Fig. 3A, S4, S5). The set of retrained models that make use of the standard curves overall performs best across the validation datasets, at least when the mean absolute error and correlations between experimental and predicted scores for PTEN are not considered (Fig. 3A, S4, S5). Interestingly, this set of retrained models also captures the ranking of variant effects, a measure that does not depend on whether standard curves are used for validation or not, better than models trained without the standard curves for several proteins, including PTEN (Fig. 3A). Together, these results show that it is possible to learn variant effects across several experimental datasets and that it is advantageous to model the mapping from low- to high-throughput score space during training.

**Figure 3.**
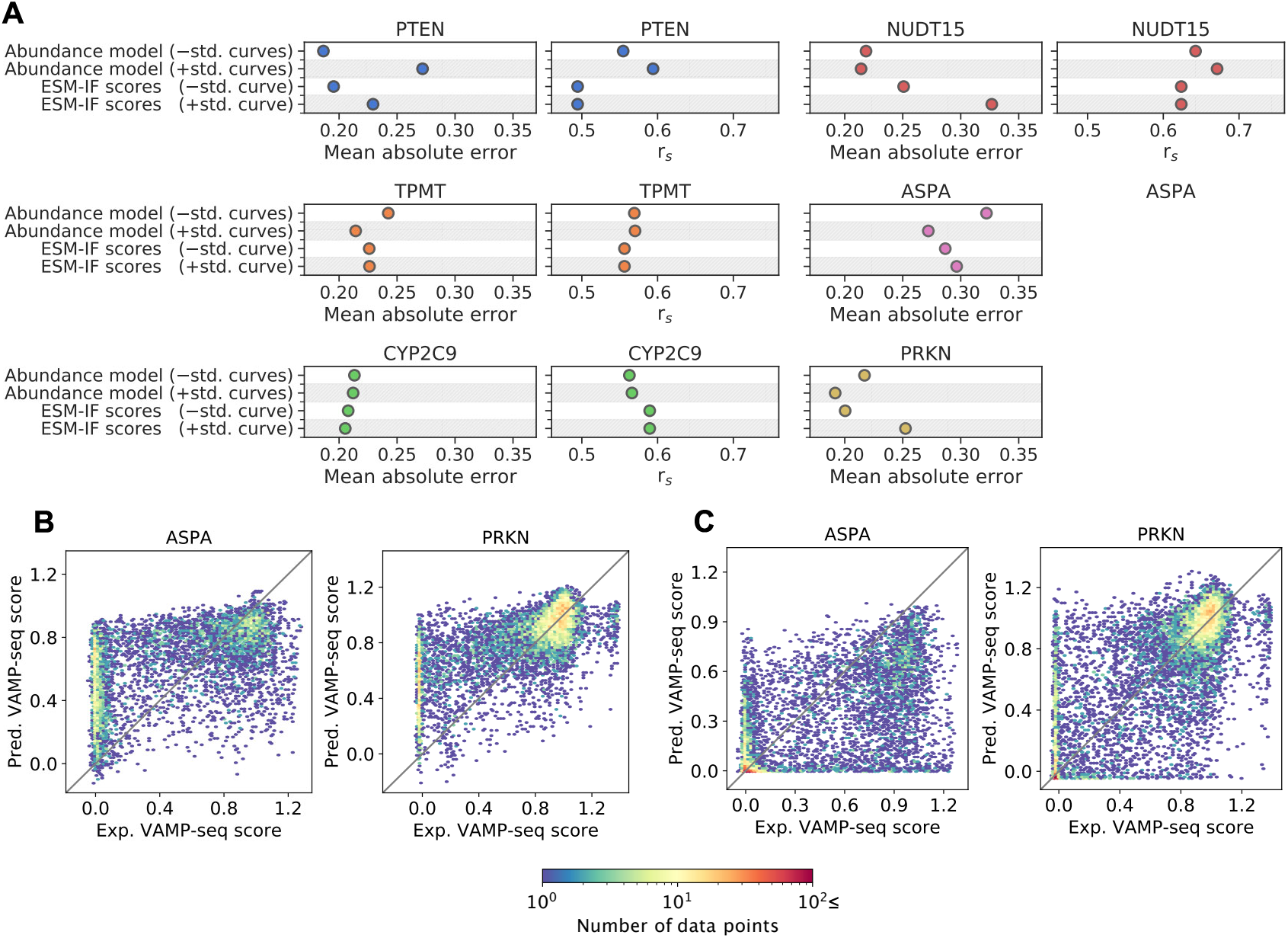
Results from leave-one-protein-out cross-validation of learning framework trained without or with experimentally informed standard curves downstream of the abundance network and benchmarked against zero-shot ESM-IF predictions of VAMP-seq scores. (A) Mean absolute error and *r_s_* between experimental and predicted VAMP-seq scores from abundance models trained without (−std. curves) or with (+std. curves) standard curves. Results are shown for all six proteins, and predictions for each protein were obtained with models trained without seeing data for that protein. Here, we only show results from models trained without PTEN VAMP-seq data, but results from runs that include this data in the training set can be found in Fig. S4, S5. The mean absolute error and *r_s_* between VAMP-seq scores and zero-shot ESM-IF scores are also shown. The ESM-IF scores were either compared directly to VAMP-seq scores during benchmarking (−std. curve) or mapped to the low-throughput score space and run through the standard curves before comparison to the experimental data (+std. curve). Generally, areas marked in grey indicate that the standard curves were used for training and/or validation. (B) Example scatter plots of experimental and predicted VAMP-seq scores for ASPA and PRKN from abundance models trained *without* standard curves (i.e. Abundance model (−std. curves)). (C) Example scatter plots of experimental and predicted VAMP-seq scores for ASPA and PRKN from abundance models trained *with* standard curves (i.e. Abundance model (+std. curves)). Similar plots can be found for all proteins in Fig. S6.

**Figure 4.**
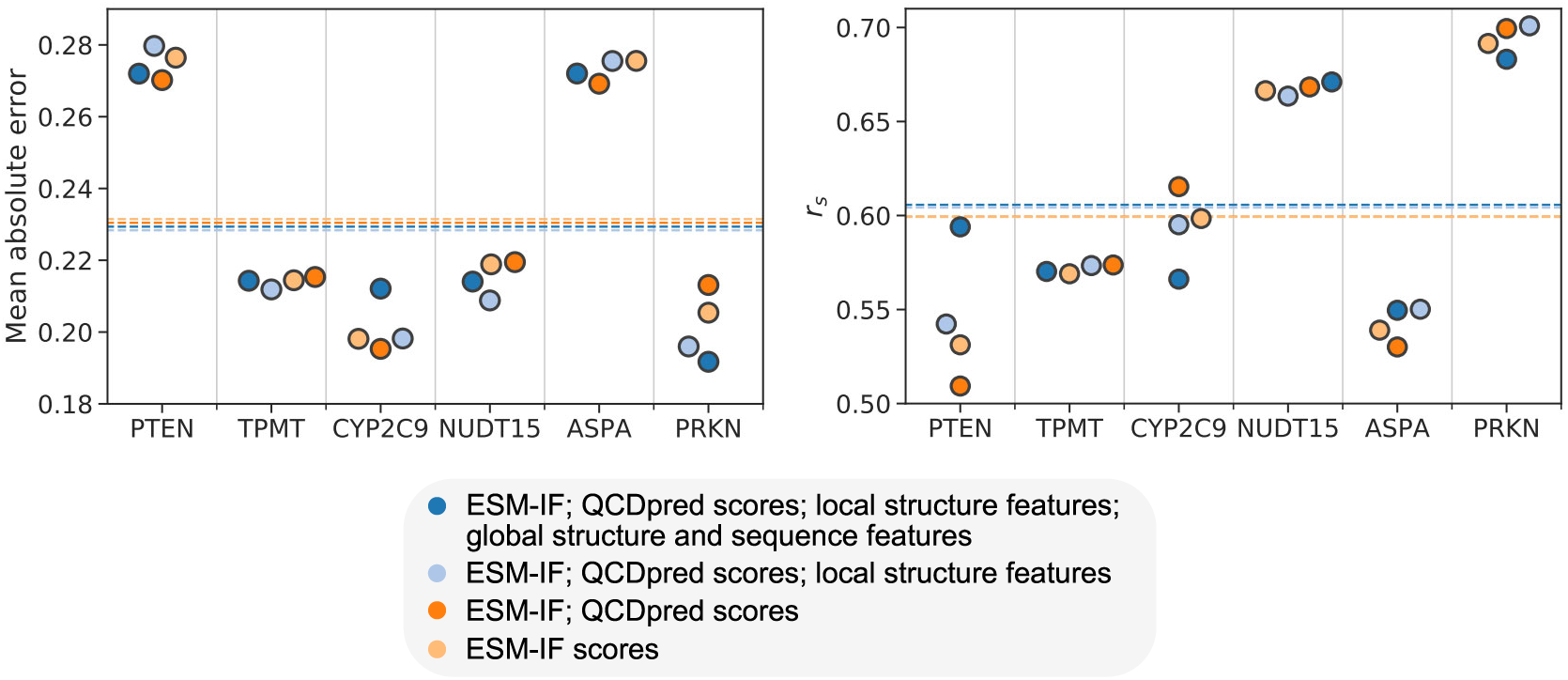
Handcrafted features have minimal influence on model performance. The importance of handcrafted features (that is, all features aside from ESM-IF scores) was studied by training abundance models (including standard curves) in a leave-one-protein-out cross-validation fashion using different sets of input features. The input feature sets range from including all handcrafted features (dark blue) to including only ESM-IF scores (light orange). A detailed overview of features in the four feature sets can be found in Fig. S2. The mean absolute error and *r_s_* between experimental VAMP-seq scores and scores predicted using the different feature sets are shown for the individual (validation) proteins. Dashed lines indicate the average mean absolute error or *r_s_* across proteins.

We compared the scatter plots of experimental against predicted abundance scores for both sets of models (Fig. 3B,C, S6). When the abundance network is trained and validated *without* passing the network output through the standard curves, VAMP-seq scores for variants with WT-like abundance appear relatively well-predicted. However, variants of NUDT15, ASPA and PRKN with experimental VAMP-seq scores close to 0 are often predicted to have abundance scores higher or much higher than the experimental values (Fig. 3B, S6A). The low- to high-throughput score mappings for these three proteins (Fig. 1A) suggest that a considerable number of these variants would, in fact, *not* have nonsense-like abundance in a low-throughput experiment. Since this information is not intrinsically present in the high-throughput scores or in the learning framework that does not include the standard curves, the abundance of these variants cannot be correctly predicted. As expected, predictions from the learning framework *with* standard curves are more accurate for ASPA and PRKN variants with high-throughput abundance scores of or close to 0 (Fig. 3C, S6B). However, variants that were previously correctly predicted to have wild-type-like abundance scores or at least not nonsense-like abundance scores are now assigned scores that are too small, suggesting that it might be possible to improve the model accuracy by using an fine-tuned set of standard curves rather than our initial fits. We have experimented with such fine-tuning and report the results of this in the supplementary material.

### Supervised abundance predictor outperforms baselines for some proteins

We next evaluated the performance of our models by benchmarking against three different baselines, specifically (*i*) VAMP-seq score predictions from a set of structure context-dependent substitution matrices (Schulze and Lindorff-Larsen, 2024), (*ii*) Rosetta predictions of changes in folding free energies (ΔG = ΔG_variant_ − ΔG_wild type_) (Park et al., 2016) and (*iii*) zero-shot predictions from ESM-IF corresponding to the ESM-IF scores that we aim to improve upon by training against the VAMP-seq data. Below, we focus on the benchmarking against the ESM-IF scores, since we found ESM-IF to be the best-performing baseline model and moreover use the model upstream of the abundance network in our learning framework (Fig. 2).

Since our goal was ultimately to develop an abundance model for predicting variant effects as measured in low-throughput experiments, the accuracy with which our models and the baseline models predicts results from low-throughput experiments would be relevant to evaluate. However, since only few low-throughput measurements have been reported for each of the proteins in our dataset, evaluation on these data gives relatively limited insight into the utility of the individual models (Fig. S7). For a more robust assessment, we therefore mainly evaluated the models using the high-throughput scores and assumed that if a model captures variant effects in the high-throughput space better than another model, the first model should also be the more accurate low-throughput score predictor. Since we aimed to learn to predict scores correctly on an absolute scale, we did not only compare the *r_s_* between experimental and predicted scores across supervised and baseline models, but also focused on comparing their mean absolute errors.

We took two separate approaches to evaluate absolute errors of baseline model predictions in the high-throughput score space. We first considered how well the baseline models predict high-throughput scores when information about the mappings from low- to high-throughput data is not provided, that is, when predictions are directly compared to the high-throughput data. To facilitate a more fair comparison with our trained models, we also mapped baseline model predictions to the low-throughput score space, passed the mapped scores through the standard curves and finally compared the standard curve output to the experimental data (Fig. S8, S9, S10). The details and results of both approaches are discussed in the supporting information. Below, we thus compare the performance of our trained models to that of ESM-IF zero-shot predictions in their raw and standard-curve transformed forms (Fig. 3A).

We found that our abundance model (incl. standard curves) trained against the VAMP-seq data on average slightly improves variant effect ranking; the average *r_s_* across proteins is 0.59 for the ESM-IF baseline and 0.61 for the trained model. The average mean absolute error is 0.23 for both our trained model and the baseline predictions that are not standard-curve transformed. Standard-curve transformation of the baseline predictions increases the average mean absolute error to 0.26 (Fig. S11, S12). Whether the trained abundance model is more accurate than the ESM-IF baseline or not depends on the validation protein (Fig. 3A); for TPMT, NUDT15, ASPA and PRKN, our abundance model trained with standard curves downstream of the abundance network predicts absolute VAMP-seq scores most accurately (that is, with the lowest mean absolute error), and for PTEN, TPMT and NUDT15, the supervised model is better than the baseline when it comes to ranking variant effects (that is, has the highest *r_s_*).

We moreover observe that sometimes, for example in the case of CYP2C9, ESM-IF zero-shot predictions are the most accurate across several evaluation metrics, indicating that the supervised ‘learning’ can make predictions noisier than the model input features originally were. In some cases, the supervised training performed without standard curves downstream of the abundance network results in worsening of prediction accuracy compared to the ESM-IF baseline; when the standard curves are used during training, this worsening can be rescued, as seen for example for TPMT, ASPA and PRKN. Finally, it is also possible to improve over the baseline when the framework is trained without standard curves, as we have shown for PTEN and NUDT15 (Fig. 3A).

Overall, our results show that it is possible, but challenging, to supervise a model to improve over the ESM-IF baseline when training and validating across VAMP-seq datasets. Importantly, we show that explicitly accounting for the influence of the experimental process on the VAMP-seq scores is useful and even necessary to compete with or improve over the baseline, emphasising the general importance of modelling the experimental process as part of across-dataset learning approaches (Fig. S11, S12). We note that while our supervised abundance model and ESM-IF on average perform approximately similarly in terms of VAMP-seq score predictions, only our model (or, specifically, the abundance network) is trained to output scores on the absolute scale of the low-throughput scores (Fig. S7). The improved VAMP-seq score prediction accuracy of our learned model compared to that of the standard curve-transformed ESM-IF scores shows that the abundance network works better as a low-throughput score predictor on an absolute scale than ESM-IF predictions linearly mapped to this scale (Fig. 3A, S9). Our supervised learning framework has thus allowed us to learn a low-throughput score predictor although the experimental information on this score space is sparse.

Finally, we emphasise that our way of benchmarking the supervised abundance model against baseline model predictions, including ESM-IF scores, is biased towards favouring the baselines when mean absolute errors are compared. The baseline model predictions are linearly mapped to either the high- or low-throughput score space on a per protein-basis using all available experimental data for the individual systems (Fig. S8, S9, S10), meaning that the absolute scales of the baseline model scores are adjusted *per protein* using experimental data for that protein. In contrast, the supervised model needed to learn to predict scores on the correct absolute scales across proteins to perform well and was not calibrated on the experimental data that it was validated on. In other words, we have set up a strict benchmarking when it comes to the absolute errors, but in spite of this, the supervised model still performs as well as, and in several cases better than, all of the baselines (Fig. 3A, S11, S12). An important advantage of our model is thus that given a new protein, it will output variant effect scores on an absolute and interpretable scale.

### Analysis of abundance network input features

The results presented so far were generated using all of the stability- and cellular degradation-related properties listed in Fig. S2 as abundance network input features. The features were se-lected to provide information about the local and (sub)global environment of the variant residue site. Since our abundance model was tasked with learning variant abundance effects on an absolute scale, we hypothesised that the model would benefit from input features containing information related to absolute local and global folding stability. We tried to model these stabilities by using relatively simple structural features, such as residue solvent accessibility, packing density and depth, as well as AlphaFold2 pLDDT scores and predicted, domain-level absolute folding free energies (Cagiada et al., 2025). We moreover used QCDpred (Johansson et al., 2023b) to predict the quality control degron potential of all 17-residue sequence segments in the six proteins.

However, not all of these features are necessarily useful or relevant to predict substitution effects on abundance. We therefore next analysed the importance of different feature groups and tested if our handcrafted features added useful information to the prediction problem. To perform this analysis, we retrained and revalidated our standard-curve-based learning framework with leave-one-protein-out cross-validation three times, for each retraining reducing the number of abundance network input features. We first removed features related to the absolute folding stability and quality control degron potential of each protein or domain for multidomain proteins. We then further removed all features related to the local structural environment of the target residue site. Finally, we removed all input features except for wild-type and variant residue one-hot encodings and the 20 ESM-IF scores that represent the effects of having each possible standard amino acid at the target residue site (Fig. S2).

The differences between using the full and reduced sets of features as input to the abundance network are generally small (Fig. 4). The average model performance across proteins is equally good for the four feature sets, and changing the input feature set mostly does not affect predictive performance for individual proteins. For selected proteins, like PTEN, CYP2C9 and PRKN, the choice of feature set does seem to have some influence on the accuracy of predictions. We note, however, that the ranking of feature sets in terms of mean absolute error or correlation between predicted and experimental VAMP-seq scores depends on the validation protein. For example, the best feature set for PTEN predictions is the full feature set, but this feature set results in the worst predictions for CYP2C9, at least when performance is evaluated using *r_s_*. Based on this, we con-clude that some of our handcrafted features likely provide useful information in addition to the ESM-IF scores. On average, however, our handcrafted features do not contain more information than the ESM-IF scores alone, and it thus seems like we have not managed to construct a set of features that are universally informative. We note that we have very few training examples for features that are at the protein or domain level.

We have shown that predicted ΔΔG values and ESM-IF scores capture abundance effects of many variants, but they clearly do not capture all variation in the VAMP-seq data, regardless of the non-linear relations between variant abundance scores from low- and high-throughput experiments. Although ESM-IF scores and ΔΔG values correlate (Fig. S13) (Hsu et al., 2022b; Reeves and Kalyaanamoorthy, 2024), ESM-IF generally captures the impact of substitutions on abundance better than the ΔΔG values calculated with Rosetta (Fig. S12, S14), with effects of substitutions of solvent-exposed residues especially described better by ESM-IF scores (Fig. S14). We have previously shown that QCDpred (Johansson et al., 2023b), our sequence-based predictor of quality control degrons, can capture substitution effects on abundance in exposed regions of PRKN (Clausen et al., 2024). Substitutions that increase or decrease abundance through modulation of quality control degrons are not well-described by Rosetta and ESM-IF (Fig. S15), but can be described with QCDpred in PRKN as well as in ASPA (Fig. S15), highlighting concrete types of variant effects that features beyond ESM-IF scores are relevant for.

## Conclusions

Building on our previous work (Schulze and Lindorff-Larsen, 2024), we here trained a variant abundance predictor using ca. 32,000 variant effect scores from VAMP-seq experiments, a MAVE method that estimates the effects of residue substitutions on the steady-state abundance of proteins in human cells. VAMP-seq quantifies variant effects in high throughput through a combination of fluorescence-based sorting and sequencing of variant-expressing cells, an approach that might lead to non-linear and experiment-specific mappings from molecular level-variant effects to high-throughput readout.

We compared low-throughput measurements of variant abundance to VAMP-seq scores from six independent experiments and indeed found that the mapping between the two data types varied across experiments. This observation suggested that supervised training of an abundance predictor across independent datasets might benefit from dataset-specific modelling of the influence of the experimental method on the high-throughput readout. Generally, this observation also highlights that it is non-trivial to aggregate data across large-scale mutagenesis datasets, even if molecular phenotype and experimental technique are kept constant.

As a result, we developed a supervised learning framework that allowed variant effects to be learned across datasets, specifically by incorporating a shared variant effect prediction module and a set of dataset-specific standard curves that accounted for the mapping from variant abundance to VAMP-seq scores for individual datasets. We applied the framework to a collection of six VAMP-seq datasets to train a variant abundance model and found that the resulting model outperforms a predictor trained without standard curves. While our supervised abundance model overall has a performance comparable to that of the baseline model ESM-IF, we stress that our model was tasked with predicting variant effects on an interpretable scale, specifically on the scale of low-throughput measurements of WT-normalised cellular fluorescence intensities, a challenging target to learn across proteins.

We hypothesise that our approach is currently limited, at least in part, by inaccuracies in our representations of the standard curves and that a refinement of these would enable improved abundance model accuracy. Our work highlights the importance of collecting high-quality low-throughput data to validate, interpret and model MAVE results. In relation to that, we note that experiment-specific and non-linear relations between an underlying molecular phenotype or low-throughput validation scores and high-throughput MAVE data is not exclusively a VAMP-seq-specific phenomenon (Tareen et al., 2022; Gersing et al., 2024) and should generally be considered when interpreting and modelling large-scale mutagenesis datasets. The basic idea of our framework, namely to train a shared variant effect predictor with the help of downstream modules to account for training data differences, might thus be relevant to apply more generally to MAVEs that are not VAMP-seqs.

We also expect future abundance model improvements to design and include features that describe the tendency of proteins and their variants to become degraded in cells. To fully model the effect of a substitution on cellular protein levels, it might be necessary to describe how the equilibrium (and potentially kinetics) between the wild-type and degradation-prone states of a protein is affected by the substitution and how different components of the cellular degradation machinery interact with both of these states, potentially leading to complicated kinetic models of proteostasis (Bershtein et al., 2013; Powers and Gierasch, 2021). In our modelling framework, the abundance network is tasked with learning the relation between a range of stability- and degradation-related features and cellular protein levels. As an alternative to this, one might try to describe the relation between the input features and abundance using biophysical modelling to create a more mechanistic and interpretable model (Bershtein et al., 2013), for example by assuming a sigmoidal relationship between substitution impact and abundance (Sarkisyan et al., 2016; Johansson et al., 2023a; Faure et al., 2022).

Our work here is related to other approaches to account for the influence of the experimental process on MAVE results (Otwinowski and Nemenman, 2013; Tareen et al., 2022; Johansson et al., 2023a; Faure and Lehner, 2024). Since MAVE measurement processes might introduce experiment-specific and non-linear relationships between variant phenotype and MAVE readout, the readout does not necessarily provide a directly interpretable quantification of variant phenotypes. Several approaches exist to infer underlying variant phenotypes directly from high-throughput data, for example by assuming a linear relationship between phenotype and MAVE readout (Weng et al., 2024; Faure and Lehner, 2024) or by using multi-mutant information to fit non-linear relationships (Tareen et al., 2022; Johansson et al., 2023a). Our approach here is different, since we make use of available low-throughput measurements to describe how the measurement process affects the high-throughput readout. As a result, the function to describe the measurement process does not need to be assumed, and the approach should limit the potential of the experiment modelling to compensate for flaws in the mapping from genotype to variant effect that is simultaneously being learned.

## Methods

### Preparation of VAMP-seq data and low-throughput luorescence intensity data

We collected VAMP-seq scores and WT-normalised GFP:mCherry fluorescence intensity ratios from low-throughput experiments from previously published work (Matreyek et al., 2018, 2021; Amorosi et al., 2021; Suiter et al., 2020; Grønbæk-Thygesen et al., 2024; Clausen et al., 2024). We used the most recently reported PTEN VAMP-seq dataset, which combines data from two independent VAMP-seq experiments to maximise the mutational completeness of the dataset (Matreyek et al., 2018, 2021). We excluded VAMP-seq scores from the N-terminal transmembrane helix of CYP2C9 (corresponding to the first 28 residues) to only study variant effects in soluble parts of the proteins. We generally excluded VAMP-seq scores for synonymous variants and nonsense variants to make our combined VAMP-seq score dataset consist only of single residue substitution effect scores. However, as low-throughput experiments were conducted for nonsense variants and synonymous variants of some of the proteins in our dataset, we did include both low- and high-throughput scores for these variants to fit the standard curves shown in Fig. 1. Low-throughput measurements of GFP:mCherry fluorescence intensity ratios were, if possible, collected from the supporting material of the individual VAMP-seq papers or otherwise extracted directly from reported plots. Measurements were normalised to the values reported for wild-type proteins in cases where such normalisation had not already been performed in the original work.

### Structure preparation and calculation of structure-based features

We previously generated a set of protein structures combining crystal structure information with AlphaFold2 (Jumper et al., 2021) and ColabFold (Mirdita et al., 2022) structure predictions to analyse substitution effects on abundance in the six VAMP-seq proteins (Schulze and Lindorff-Larsen, 2024). The idea behind combining structural information from several sources was to stay as close to the crystal structure conformations as possible and at the same time have structural information for residues for which no crystal structure data was available. We reused our combined experimental and predicted structures in this work. The structures are all wild-type structures, meaning that none of our analyses make use of variant structure data. As in our previous work, we used NUDT15 and ASPA homodimer structures as input for all analyses (Suiter et al., 2020; Grønbæk-Thygesen et al., 2024; Schulze and Lindorff-Larsen, 2024). Structure visualisations were created with ChimeraX (Pettersen et al., 2021).

Using this set of input structures, we derived a number of structure-based features. We calculated absolute solvent accessible surface areas for all residues with DSSP (Kabsch and Sander, 1983) and normalised the absolute values to the theoretical maximum solvent accessibility for every amino acid residue type (Tien et al., 2013) to obtain relative accessibilities (rASA). We also used DSSP to calculate hydrogen-bonding energies for all hydrogen-bonding backbone atoms and added the energies up for each residue to estimate the total backbone hydrogen-bonding energy per residue. We calculated the average residue depth (that is, the average structural depth of the atoms that make up a given residue) for all residues in the six structures using the DEPTH web-server (Chakravarty and Varadarajan, 1999; Tan et al., 2013). A weighted contact number was also calculated for all residues as previously described (Schulze and Lindorff-Larsen, 2024). AlphaFold2 pLDDT scores were obtained from structures in the AlphaFold2 structure database (Varadi et al., 2022). Rosetta ΔΔG values (ΔΔG = ΔG_variant_ − ΔG_wild type_) were estimated with the cartesian_ddg protocol and *opt-nov15* Rosetta energy function (Park et al., 2016) and taken from our previous work (Schulze and Lindorff-Larsen, 2024).

### Calculation of ESM-IF scores and predictions of absolute folding stabilities

We used the structure-based ESM-IF model (Hsu et al., 2022b) to evaluate substitution effects of the target proteins. Starting from the pre-trained ESM-IF model (esm_if1_gvp4_t16_142M_UR50, https://github.com/facebookresearch/esm), we input the structure and sequence of the target protein and used the ESM-IF language model decoder to calculate conditional likelihoods for all possible substitutions at each residue position (that is, the likelihoods for all possible single substitution variant sequences given the wild-type protein structures). As in previous work (Hsu et al., 2022b), we then used these likelihoods to calculate the ratio between the variant and wild-type sequence likelihoods for all variants. We report the logarithm of these likelihood ratios in this paper and refer to these log-transformed likelihood ratios as ESM-IF scores.

We predicted the folding free energy (ΔG_f_) per protein or per domain for multidomain proteins using our previously published ESM-IF-based prediction method (Cagiada et al., 2025). The method predicts ΔG_f_ for small, single-domain proteins by summing up the predicted likelihoods of finding the wild-type sequence amino acid residue at each residue position in the structure. We estimated ΔG_f_ for domains in multidomain proteins in the context of the full protein by running ESM-IF on the full protein and summing up the wild-type residue likelihoods for residues on a per-domain basis. Our approach assumes that folding of individual domains is stabilised by interactions with the rest of the protein and that individual domains unfold independently.

### Calculation of quality control degron scores with QCDpred

QCDpred is a logistic regression model trained to predict whether a given 17-residue peptide is likely to act as a quality control degron in yeast cells (Johansson et al., 2023b) by causing cellular degradation of a reporter protein that is covalently bound to the peptide (Mashahreh et al., 2022). We used QCDpred to predict quality control degron scores for all residues in the wild-type and variant protein sequences of the six proteins in our dataset. We ran the predictor on all possible 17-residue windows, or tiles, of the six wild-type sequences. We then assigned the QCDpred score for each 17-residue tile to the central residue of the tile (that is, residue 9) to thereby obtain a quality control degron score per wild-type residue. In each tile, we moreover substituted the central residue to the remaining 19 possible residues and calculated a new QCDpred score per substitution. This gave us degron scores for the variants. QCDpred scores for N-terminal (or C-terminal) residues were calculated by gradually decreasing the sequence window size so that the number of residues towards the N-terminal (or C-terminal) of the ‘central’ residue was reduced below 8, while the number of residues towards the C-terminal (or N-terminal) of the ‘central’ residue was fixed to 8. QCDpred scores for terminal residues were thus calculated using tiles shorter than 17 residues, and we corrected for changes in the tile size *n* by multiplying the sum over logistic regression parameters with a factor 17/*n*.

We moreover estimated a degradation score per protein or per domain in multidomain proteins by taking the average QCDpred score over residues belonging to that domain or protein, similarly to what was done for disordered regions in previous work, and which was shown to correlate with cellular life times of proteins (Johansson et al., 2023b). Since the QCDpred score for a residue contains information about the types of residues sitting -8 and +8 residues relative to it, scores for residues positioned close to domain boundaries will be somewhat influenced by residues in the neighbouring domain, but we assume that the contribution from neighbouring domains is negligible.

### Domain segmentation

We split proteins into domains according to a combination of the Representative Domains entry in InterPro (Paysan-Lafosse et al., 2022) and visual inspection of the protein structures, except for PRKN, which we split following a previously published PRKN domain overview (Seirafi et al., 2015). According to InterPro, TPMT, CYP2C9, NUDT15 and ASPA monomers each consist of only a single domain, while PTEN consists of two domains. Across the proteins, several residues do not belong to any of the domains specified in InterPro. These residues are often predicted by AlphaFold2 to have low pLDDT scores and are found in both terminal and internal regions of the proteins. In cases where such residues make up a continuous stretch of 25 residues or more, we grouped the residues to form a pseudo-domain. In all other cases, we assigned such residues to belong to the neighbouring domain(s). The domain segmentation was only relevant for calculation of per-domain ΔG_f_ values and average QCDpred scores; all other structure- and sequence-based features were derived from full-length structures and sequences.

### Fitting of standard curves

Standard curves mapping low-throughput measurements of fluorescence intensity ratios to VAMP-seq abundance scores were fit with the MLPRegressor class from scikit-learn (v1.0.2). All standard curves were fit using a single hidden layer consisting of two nodes, hyperbolic tangent activation functions, the lbfgs solver, L2 regularisation with alpha set to 0.001 and a maximum of 500 iterations. All other parameters were set to the class defaults, except the random_state parameter. One MLP-based standard curve was determined per VAMP-seq dataset, and all MLPs were fit with a single input and a single output node, corresponding to an input low-throughput fluorescence intensity ratio and an output VAMP-seq score for a single variant.

To obtain the final set of standard curves reported in Fig. 1, the standard curves were for every protein fitted up to 20 independent times using for each fit a unique random_state parameter value from the integer interval from 0 to 20 for initialisation of MLP weights and biases. We required that all standard curves should be monotonically increasing in the low-throughput score range from -1 to 2 and not output values smaller than -2 for an input value of -1. The standard curve fits that passed these filters did for the individual proteins not display any notable differences in the experimentally informed input value, or fluorescence intensity ratio, range. The standard curve that for each protein was fitted with the smallest random_state integer value that allowed the curve to pass the described filters was selected as the final standard curve for that protein and used for all subsequent analyses, and the standard curves selected this way are shown in Fig. 1.

### Model architecture and training protocol

Our model architecture for learning variant effects across VAMP-seq datasets consisted of (*1*) ESM-IF to generate features for downstream modelling, (*2*) a downstream dense neural network that we refer to as the abundance network and (*3*) a module consisting of six dataset-specific functions or standard curves that map the output of the abundance network to a predicted score in VAMP-seq score space (Fig. 2). We used supervised learning against multiple VAMP-seq datasets to parameterise the abundance network and in a few cases also the standard curves. We performed model training several times using various sets of training data, input features and hyperparameters.

Generally, the abundance (dense neural) network consisted of an input layer of between 60 and 69 nodes depending on the number of input features used (see Fig. S16) and was trained to output a single number. The network moreover had one or two hidden and fully-connected layers with a variable number of nodes in each layer; the specific numbers of hidden layers and nodes were determined by hyperparameter tuning of the individual models and are specified in Table 1. We used the leaky ReLU activation function (He et al., 2015) and applied batch normalisation (Ioffe and Szegedy, 2015) between all abundance network layers. The individual standard curves in the standard curve module were also modelled as small networks that each consisted of a two-node hidden layer with one-node input and output layers. We used the hyperbolic tangent activation function between all layers in the standard curve networks. We initially fit the standard curve network for each protein with the scikit-learn MLPRegressor class (see *Fitting of standard curves*) and then used the fitted weights and biases as starting point for the abundance model training.

**Table 1.** Overview of trained models, including architecture, training data, input features and hyperparameters used for the individual models. *Model architecture*: Indicates whether modules were trained without or with standard curves downstream of the abundance network, i.e. whether the architecture in Fig. 2 or Fig. S3 was used. *Trained module(s)*: Indicates whether only abundance network parameters or both abundance network parameters and standard curve parameters were updated during training. *Training data*: Indicates whether all VAMP-seq datasets were used for leave-one-protein-out cross-validation, or if PTEN VAMP-seq data was consistently omitted from the training data during cross-validation. *Input feat.*: Specifies which input feature set was used for training. The feature set numbering follows the overview in S2. *Abundance net.*: Architecture of the abundance network, with (x,y) specifying a network with x nodes in the first hidden layer and y nodes in the second hidden layer *LR*: Learning rate. *BS*: Batch size.

| Model type | Trained module(s) | Training data | Input feat. set | Abundance net. | LR | BS |
| --- | --- | --- | --- | --- | --- | --- |
| –std. curves | Abundance net. | All | 1 | (64,0) | 0.05 | 16 |
| –std. curves | Abundance net. | –PTEN | 1 | (8,0) | 0.05 | 128 |
| +std. curves | Abundance net. | All | 1 | (64,0) | 0.05 | 16 |
| +std. curves | Abundance net. | –PTEN | 1 | (64,32) | 0.01 | 128 |
| +std. curves | Abundance net. | –PTEN | 2 | (16,8) | 0.05 | 32 |
| +std. curves | Abundance net. | –PTEN | 3 | (64,0) | 0.005 | 64 |
| +std. curves | Abundance net. | –PTEN | 4 | (64,0) | 0.005 | 32 |
| +std. curves | Abundance net. + std. curves | All | 1 | (8,0) | 0.001 | 16 |

Model training was performed using PyTorch (v1.2.0) (Paszke et al., 2019). We parameterised all models through minimisation of the mean absolute error between model output and experimental VAMP-seq scores. Minimisation was performed using Adam (Kingma and Ba, 2017) with learning rates and batch sizes as noted in Table 1.

Models were always trained using leave-one-protein-out cross-validation, meaning that all data for five proteins was used as training data, while data from the sixth protein was used for validation. VAMP-seq data for variants that were used to fit standard curves was always excluded from the validation data, but not from the training data. For every training-validation split, we always trained an ensemble of three models, initialising weights differently in the individual rounds of training. Training was performed using early stopping with a patience of three, with patience based on the mean absolute error to the validation data. Final variant effect scores reported for validation proteins were calculated by averaging predictions from the three model-ensembles. We generally report predictions for all variants of the validation protein, except we always leave out variants that were used for fitting of standard curves. This is also the case for models trained without standard curves and for baseline models.

### Hyperparameter tuning, feature selection and feature normalisation

We report leave-one-protein-out cross-validation results for several models that differ in overall model design, for example due to variation in the application of standard curves or in the used input features and hyperparameters. A detailed description of the in total eight different model designs are outlined in Table 1. For each design, the abundance network size, learning rate and batch size were optimised based on a random search over the parameter space defined in Fig. S16. For each of the eight designs, we specifically performed leave-one-protein-out cross-validation using 50 random combinations of the three hyperparameters (randomly sampling combinations from the possible hyperparameter values listed in Fig. S16), and picked the hyperparameter combination that resulted in the smallest average mean absolute error across validation proteins. In the one model design where standard curves were reparameterised during training, we picked the hyperparameter set that simultaneously minimised the average mean absolute error across validation proteins and the deviation between initial and fine-tuned standard curves (Fig. S17). For designs where PTEN was excluded from the training data, we performed leave-one-protein-out cross-validation for the five remaining proteins and optimised hyperparameters based on results for those five proteins. We then trained models with the optimised hyperparameters on data for all proteins except PTEN and used the trained models to predict PTEN data.

For studies of feature importance, we trained models with subsets of the full feature set according to the feature groups defined in Fig. S2. Generally, we min-max normalised selected input features across variants in the training dataset (that is, across training proteins), and we applied the minimum and maximum feature values in the training data to also normalise input features for variants in the validation data. We specifically applied this normalisation approach to the ESM-IF scores, the weighted contact number per residue, the residue depth, the total hydrogen bonding energy per residue and the predicted ΔG_f_ per protein or domain. AlphaFold2 pLDDT scores were not min-max normalised, but simply divided by 100.

### Model training including standard curve fine-tuning

Model training that included standard curve fine-tuning was very similar to training with fixed standard curves, although with a few exceptions. As in all other cases, we started model training from an untrained abundance network and with standard curves that were fitted to the low-throughput validation data as described under *Fitting of standard curves* and shown in Fig. 1. We first trained the abundance network with frozen standard curves as described above. Afterwards, we ran another round of model training starting from the pre-trained abundance network (corresponding to the network parameterised in the previous round of training), now letting parameters of the abundance network as well as of the standard curve networks adjust. The second round of training was also performed using early stopping with a patience of three, and as before, we performed leave-out-protein-out cross-validation and trained ensembles of three models for every training-validation split.

While parameters of the abundance network were generally learned using data from all training proteins, reparameterisation of the protein-specific standard curves relied on protein-specific training data, meaning that training of the validation protein standard curve required some of the validation protein data to be used for standard curve training rather than actual validation. Thus, when standard curves were adjusted during training, we randomly split the validation protein VAMP-seq data on a variant-level and used 20% of this data to inform the fine-tuning of the validation protein standard curve. The remaining 80% of the data was never seen during training and hence used for actual validation. Importantly, the first 20% of the data was used only to inform the validation protein standard curve and never the abundance network. The validation protein data splitting was done randomly, and a new random split was used for every training run for generation of the three models in an ensemble. We compared the performance of models trained with fixed standard curves to models trained with adjustable standard curves by evaluating errors and correlation coeicients using only the 80% of validation protein variant effects scores that were selected for validation during training with adjustable standard curves.

We selected optimal hyperparameters for models with trainable standard curves by simultaneously considering (*i*) the average mean absolute error between predicted and experimental VAMP-seq scores across validation proteins and (*ii*) the distance between trained and initial standard curves. The latter consideration was necessary to not end up with standard curves that deviated considerably from the initial fits. For a given hyperparameter set and validation protein, the mean absolute error between predicted and experimental VAMP-seq scores was evaluated individually for the three independently trained base models in an ensemble and then averaged across the ensemble to estimate the prediction error for that combination of hyperparameters and validation protein. We moreover quantified the deviation between the initial standard curves and the fine-tuned standard curves by calculating the mean absolute error between VAMP-seq score predictions from both types of curves for a range of low-throughput score input values. We defined the input value range for this analysis on a per protein-basis so that the minimum and maximum values in the range corresponded to the experimental minimum and maximum low-throughput scores for each protein, and we used input scores from this range with a step size of 0.01. We averaged the mean absolute error between initial and fine-tuned standard curves across base models and validation proteins to get a final estimate of the error for a given hyperparameter set. Finally, we plotted the average mean absolute error between predicted and experimental VAMP-seq scores against the average mean absolute error between initial and optimised standard curves for all 50 hyperparameter sets (Fig. S17) and picked the hyperparameter set that best minimised both quantities.

### Benchmarking against baseline models

We benchmarked our abundance model against three different baselines: Abundance score predictions from a substitution matrix model, variant ΔΔG values calculated with Rosetta, and ESM-IF zero-shot variant effect predictions. ESM-IF variant effect predictions were obtained by calculating the ratios between variant and wild-type sequence likelihoods given wild-type protein structures as described above. Abundance score predictions from the substitution matrix model and Rosetta ΔΔG values were obtained from prior work (Schulze and Lindorff-Larsen, 2024). Briefly, we previously calculated a pair of amino acid residue substitution matrices by averaging VAMP-seq abundance scores for all possible substitution types for residues found in either solvent-exposed or structurally buried environments in the wild-type protein structures. We obtained an abundance score prediction for a given variant from these matrices by looking up the average abundance score for the variant type, using the average score for either solvent-exposed or buried residues depending on the structural environment of the target residue of the prediction. In this framework, all predictions come from leave-one-protein-out cross-validation, that is, from substitution matrices constructed by leaving out VAMP-seq data for the target protein. Buried residues were defined as residues with an rASA smaller or equal to 0.1, and exposed residues were defined as residues with an rASA larger than 0.1. Moreover, as previously described (Schulze and Lindorff-Larsen, 2024), we used the Rosetta cartesian_ddg protocol and *opt-nov15* energy function (Park et al., 2016) to estimate the difference in Gibbs folding free energy between the wild-type and variant proteins (ΔΔG = ΔG_variant_ − ΔG_wild type_) for all variants.

Since the substitution matrix model was constructed using VAMP-seq data, the abundance score predictions from this baseline model are immediately on the same scale as the VAMP-seq scores, and the mean absolute error between experimental and predicted scores can therefore easily be evaluated in a meaningful way. This is, however, not the case for the Rosetta ΔΔG values and the ESM-IF scores, which are usually found in intervals between -8 and 1 kcal/mol and between -25 and 5 (Fig. S1, S13), respectively. To calculate mean absolute errors between predictions from these baseline models and the experimental data, we performed linear regression with the VAMP-seq abundance scores as the dependent variable and either ΔΔG values or ESM-IF scores as the independent variable. We performed the regression individually for each protein and used the resulting models to transform ΔΔG values and ESM-IF scores onto the abundance score scale (Fig. S8). We then calculated the mean absolute error between these transformed values and the experimental data, thus obtaining error estimates that we could compare to the mean absolute errors between the experimental data and predictions from our abundance model.

Rosetta ΔΔG values and the ESM-IF scores also needed to be mapped to the same scale as the low-throughput measurements to be passed through the standard curves and thereby used for benchmarking that takes the standard curve transformations into account. Thus, we took an approach similar to the one just described and linearly mapped the predictions onto a the low-throughput score scale. For each protein, we fit a linear model between either ΔΔG values or ESM-IF scores and the low-throughput abundance measurements (Fig. S9), and we then used the obtained models to transform the baseline model predictions (Fig. S10). We ran the transformed values through the protein-specific standard curves to get final estimates of abundance scores in the high-throughput space and used these estimates to calculate the mean absolute error and *r* between the predictions and the experimental data.

For the leave-one-protein-out cross-validation of our abundance model without standard curves and with fixed standard curves, we report the mean absolute error and correlation coeicients between the experimental VAMP-seq data and the predictions from our model using all variant effect scores for the validation protein, except we exclude data for the few variants that were used for the initial fittig of standard curves, that is, the variants for which low-throughput data was available. To be consistent with this, the VAMP-seq abundance scores for these variants were also excluded in the analyses described in the above paragraphs and hence from the evaluation of the baseline models reported in Fig. S11, S12.

## Availability of data and code

Code and data produced in this work are available at https://github.com/KULL-Centre/_2025_Schulze_abundance-model.

## Acknowledgements

This work was funded by the Novo Nordisk Foundation challenge program PRISM (Protein Interactions and Stability in Medicine and Genomics, NNF18OC0033950, to K.L.-L.). We thank Martin Grønbæk-Thygesen, Vasileios Voutsinos, Lene Clausen, Sören von Bülow, Kristoffer E. Johansson, Amelie Stein, Rasmus Hartmann-Petersen, Douglas M. Fowler and other members of the PRISM centre for many useful suggestions and discussions about VAMP-seq and modelling of variant abundance. We acknowledge access to computational resources at the Biocomputing Core Facility at the Department of Biology, University of Copenhagen, and from the ROBUST Resource for Biomolecular Simulations (supported by the Novo Nordisk Foundation grant no. NF18OC0032608).

## Supporting information

### Fine-tuning of standard curves during abundance model training

The accuracy with which low-throughput score predictions from the abundance network are mapped to VAMP-seq scores via the standard curves influences how well our learning framework can learn and predict variant abundance effects in both low- and high-throughput space. The standard curves applied above (and shown in Fig. 1) were fit using all low-throughput score measurements available from the literature. These measurements were, however, available for only relatively few variants for each of the proteins in our dataset (on average 16.5 per protein), and the resulting set of standard curves are thus informed by these few and, in some cases, somewhat noisy data points. Related to this, the available low-throughput data does not always cover the entire relevant range of high-throughput scores. For example, no low-throughput scores were reported for variants with very low high-throughput scores for TPMT and CYP2C9. The shapes of the standard curves in these score regions are therefore somewhat arbitrary. Finally, we note that the standard curves do not always go through (1,1) even though both low- and high-throughput scores are wild-type normalised (Fig. 1).

There are thus several reasons why the standard curves might not perfectly describe the mapping from low- to high-throughput scores, and this could cause some of the problems that we observed above to be associated with using the curves. For example, the steepness of the ASPA standard curve (Fig. 1) between low-throughput score values of ca. 0.8 and 1.2 appears to be somewhat poorly described, resulting in variants predicted to have a WT-like abundance in low-throughput space to be assigned small high-throughput scores. Thus, as a way of reducing the impact of the uncertainties in the standard curve fits on the performance of our supervised model, we experimented with unfreezing the parameters of the standard curve module to let the curves adjust during model training. Specifically, we first trained the model with standard curve parameters frozen, and after convergence of training the abundance module, we unfroze the standard curve parameters and trained the standard curves and dense network simultaneously. In that way, we let errors introduced in the dense network due to uncertainties in the initial standard curves be fixed together with the standard curves.

The resulting model outperforms the model in which the curves remain frozen in terms of both absolute prediction accuracy (mean absolute error) and ranking of varinat effects (*r_s_*) (Fig. S18). However, in order to achieve this, several standard curves move relatively far away from their starting points (Fig. S19). We observe the most extreme shift in the position of the standard curve for PTEN. Training the standard curves makes several curves move further away from (1,1) than they were to begin with, suggesting that the curves become overfit. If the goal were to predict variant ranking correctly, this would not matter, since *r_s_* is improved in spite of the overfitting. However, since we aimed to predict scores correctly in the low-throughput space, it does not make sense to let the standard curves become unphysical. If the abundance network were a perfect low-throughput score predictor, the discrepancies between the predictions from the network and the high-throughput data could be used to determine the correct shape of the standard curves. However, since the abundance network is clearly not a perfect predictor, fine-tuning of the standard curves during training might in part be a way to compensate for flaws in the predictions from the network. To limit such compensation while still allowing for standard curve adjustment, the fine-tuning could potentially be performed with a strategy for regularisation of the initial standard curve fits. We did not try to implement such strategies in this work, and in the main text, we hence only show data for models trained with fixed standard curves.

Our work generally shows that the use of standard curves needs to be implemented carefully; the ability of our abundance model (trained with fixed standard curves) to learn and predict the low-throughput data and VAMP-seq scores is likely somewhat limited by imperfect descriptions of the mappings from low- to high-throughput scores. We assume that access to more low-throughput data or implementation of better strategies to refine the standard curves with existing data could facilitate training of better models than those presented here.

### Discussion of PTEN data and standard curve

The PTEN VAMP-seq dataset is relatively well-predicted by the supervised model that does not use standard curves (Fig. S5) and by all baseline models if baseline predictions are not standard curve-transformed (Fig. S11, S12). However, all model types struggle to correctly capture PTEN scores when the standard curve for the dataset is used to produce predictions (Fig. S11, S12). Across model types, the PTEN standard curve seems to introduce the problem that too many variants are predicted to have WT-like VAMP-seq scores, including variants that experimentally have VAMP-seq scores close to 0 (Fig. S6, S10). The PTEN standard curve maps a wide range of low-throughput scores between 0.5 and 1.5 to VAMP-seq scores close to 1, and small to very small low-throughput scores need to be predicted by our supervised model or the baselines to obtain VAMP-seq scores below 0.75. Clearly, our model and the baselines do not predict low-throughput scores that are small enough to transform to small VAMP-seq scores via the standard curve. To compensate for this, the PTEN standard curve moves a lot during standard curve fine-tuning (Fig. S19). The standard curve for PTEN is informed by a relatively high number of experimental data points (Fig. 1), and so it is difficult to argue that the curve should be allowed to move as much as it does in the fine-tuning experiment (Fig. S19). Thus, if the standard curve for PTEN is relatively correct, the problem would be that our model and the baselines lack information to properly describe the cellular abundance of PTEN variants; apparently, these variants are degraded to a higher degree than any of the models expect.

Clearly, more work is needed to better describe PTEN variant effects on an absolute scale (or resolve potential issues with the standard curve). Our lack of information about PTEN has implications for the performance of the supervised models that make use of the standard curves downstream the abundance network (Fig. S4, S5). To be able to evaluate the utility of a standard-curve-based learning framework, we decided to take out PTEN data from the training set for most of our trained models (Table 1). We emphasise that omitting PTEN from the training data in the cross-validation on average gives better results across the remaining proteins (Fig. S5), and that our ranking of PTEN variant effects outperforms that of all baselines by a relatively large margin when we train the abundance model with the five remaining datasets and hyperparameters that are optimised for those five datasets (Fig. S12). Accordingly, we expect a model trained without PTEN data in our current setup to generalise best.

### Detailed evaluation of baseline models

We evaluated the performance of our models by benchmarking against three different baselines. First, we compared our model predictions to predictions from a pair of residue substitution matrices that were constructed by averaging VAMP-seq scores for all possible residue substitution types separately for solvent-exposed and buried residues (***Schulze and Lindorff-Larsen, 2024***). The VAMP- seq score for a variant was predicted with these matrices by looking up the average score for the relevant substitution type, specifically using the average over either exposed or buried residues depending on the structural environment of the target residue. In our previous work (***Schulze and Lindorff-Larsen, 2024***), we used the Rosetta energy function (***Park et al., 2016***) to calculate changes in folding free energies (ΔΔG = ΔG_variant_ − ΔG_wild type_) for all variants of the six proteins. As a second baseline model, we here used these variant ΔΔG values as abundance score estimates. Finally, we also benchmarked our model against ESM-IF zero-shot predictions of VAMP-seq scores in the form of ratios of likelihoods of variant and wild-type sequences. These are the scores that we also input to the abundance network in our model and hence try to improve on by training against the VAMP-seq data.

We first considered how well the baseline models predict high-throughput scores when predictions are compared directly to the high-throughput experimental data. To evaluate the error between baseline model predictions and VAMP-seq data, we linearly mapped predictions to the VAMP-seq scale and then calculated the error between the mapped predictions and the data (Fig. S8). When evaluated this way, ESM-IF is the best variant abundance predictor of the three baseline models, although the advantage of this model over Rosetta and even the substitution matrix depends on the protein on which they are evaluated (Fig. S11, S12). This is also true when baseline models are evaluated in terms of correlations between predicted and experimental data (Fig. S12). For all three baseline models, we here and in our previous work (***Schulze and Lindorff-Larsen, 2024***) observed prediction outliers similar to those seen for our abundance model trained without standard curves, that is, high predicted scores for variants with nonsense-like abundance in the VAMP-seq experiments (Fig. S8).

These outlier patterns suggest that the predictive performance on the high-throughput data could potentially be improved by transforming the baseline model predictions via the standard curves. That would also facilitate a more fair comparison with our modelling results; such transformations would ensure that if we observe any improvement in performance for our trained models over the baselines, this would indicate that our learned model predicts low-throughput scores better on an absolute scale than the baseline models (after linear mapping to the low-throughput space), and that the prediction improvements are not only a result of the standard curve transformation that we also use during validation.

Next, we accordingly mapped predicted ΔΔG values and ESM-IF scores to the VAMP-seq score space via the standard curves. We first fit a straight line between the ΔΔG values or ESM-IF scores and all available low-throughput scores for each protein (Fig. S9A, S9B). We then transformed ΔΔG values and ESM-IF scores for all variants of each protein to predicted low-throughput score measurements via the linear fits (Fig. S10A). Finally, we ran the transformed values through the standard curves to obtain predicted VAMP-seq scores (Fig. S10B).

We observe that baseline model predictions transformed with the standard curves tend to agree less with the experimental VAMP-seq scores than the untransformed baseline model predictions (Fig. S11, S12). There might be several different reasons why this is the case. First of all, as we have already discussed, the standard curves might be overfit to the relatively few low- to high-throughput score data points that we have available (Fig. 1) and might hence not describe the relation between the two scores types entirely accurately. Moreover, the mappings from baseline model predictions to low-throughput score space are not necessarily linear (Fig. S9, S10). If our abundance model outperforms the standard curve-transformed baseline predictions, we might therefore interpret this as if we have learned a mapping from ESM-IF score to the low-throughput space that is better than the linear mapping.

---

**Figure S1.**
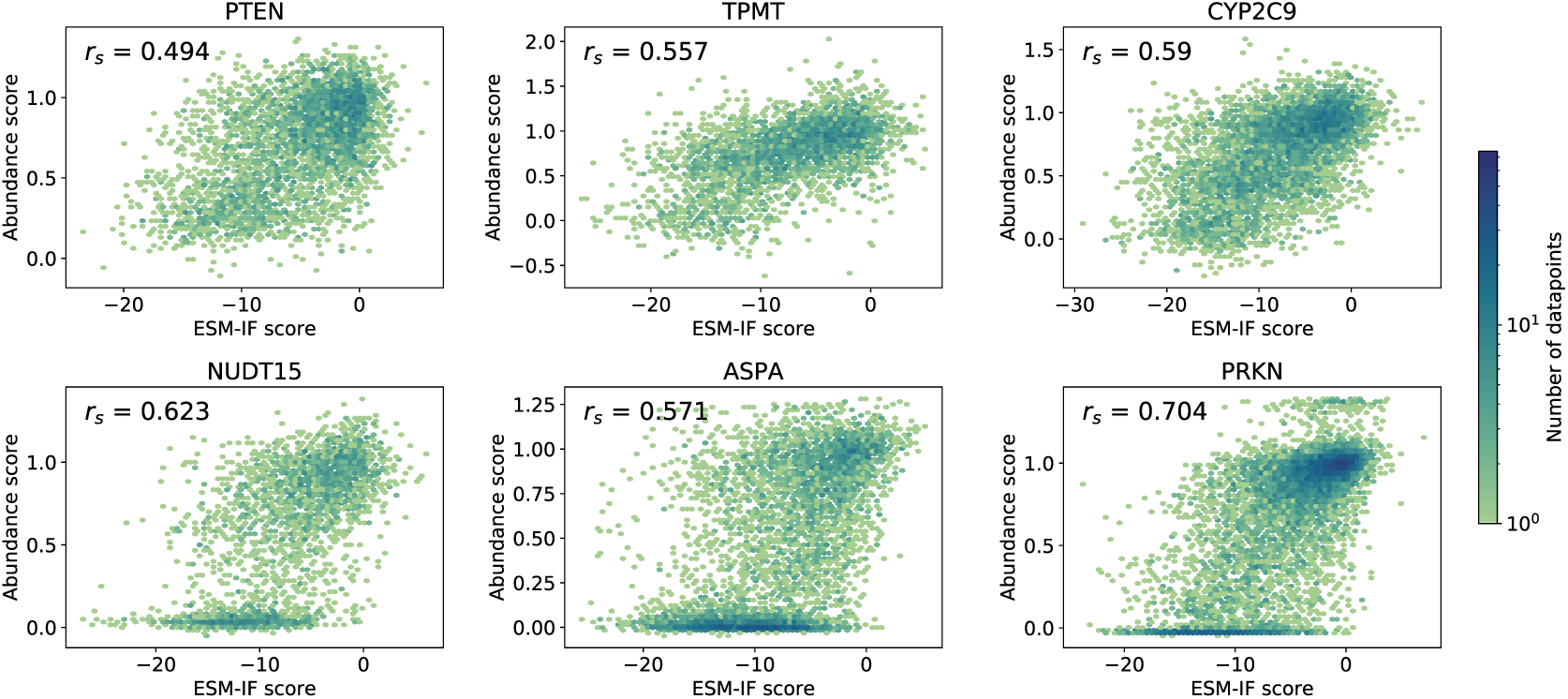
ESM-IF scores correlate with VAMP-seq scores. ESM-IF scores, specifically log-likelihood ratios of the variant and wild-type sequence likelihoods given the wild-type structure, were calculated for all single residue substitution variants of the six VAMP-seq proteins, and the correlations between ESM-IF and VAMP-seq scores are shown here for all variants of the individual proteins. The data point density is high across a range of ESM-IF scores for variants with VAMP-seq scores close to 0 for NUDT15, ASPA and PRKN, in agreement with the standard curves that for these proteins show that many variants of varying low-throughput scores were assigned high-throughput scores close to 0 in the VAMP-seq experiments. The data point density is shown with an upper limit of 75 on the colour scale.

**Figure S2.**
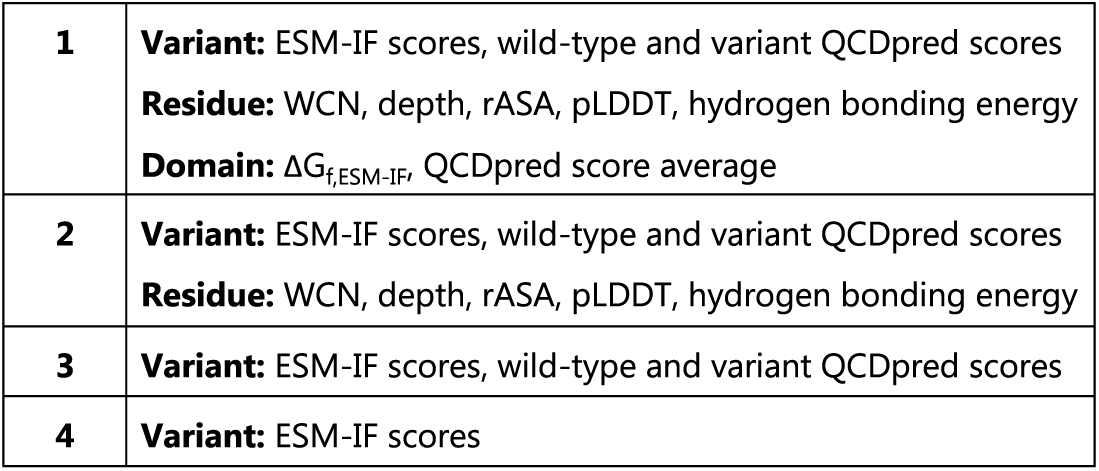
Structure- and sequence-based features used as input to the abundance network. We defined four groups of features, which we here and in Table 1 refer to with the numbers 1-4, and analysed the importance of different features for predicting variant abundance by training several abundance models that take features from different feature groups as input. As shown in Table 1, the majority of models trained in this work make use of all features, that is, feature group 1.

**Figure S3.**
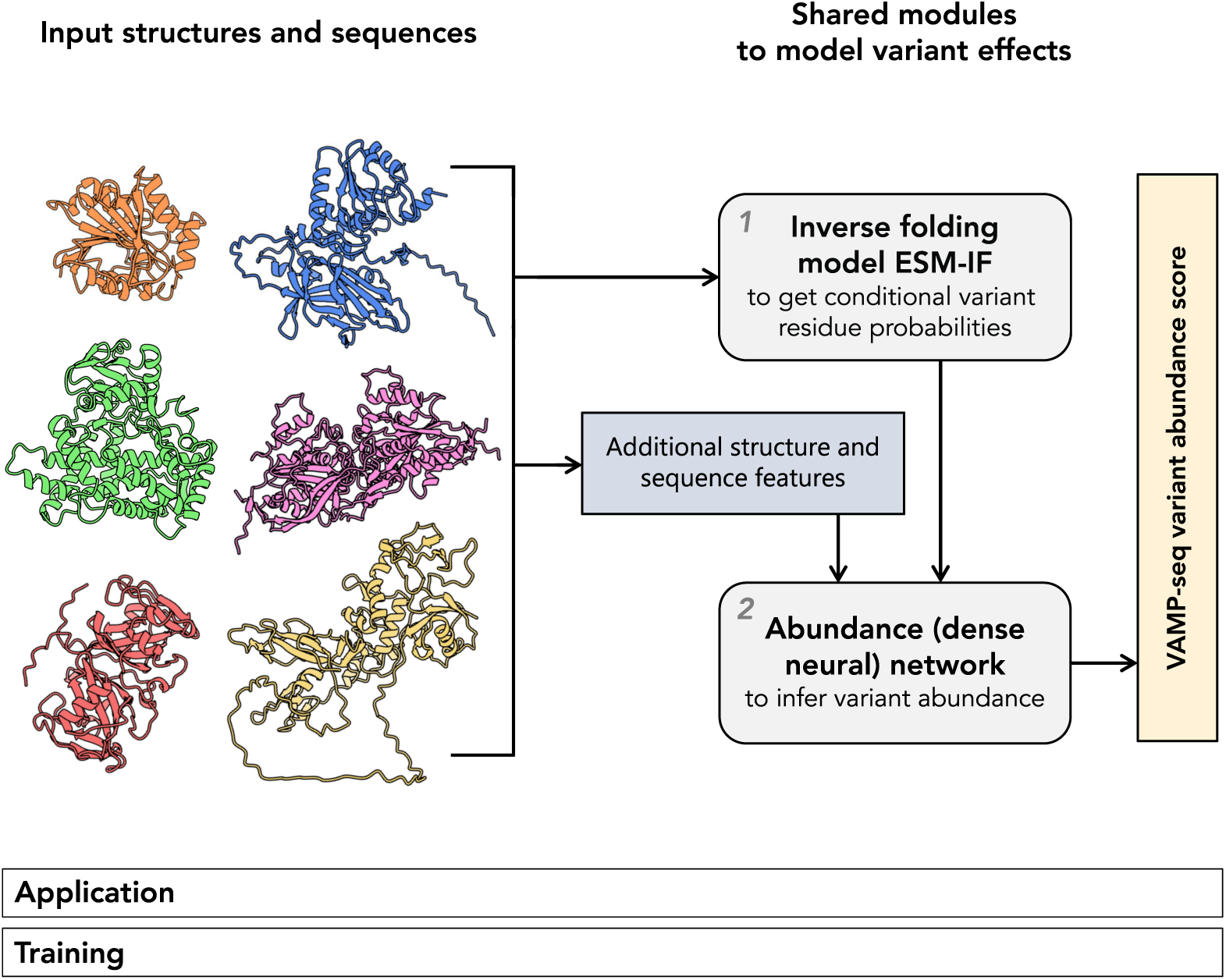
Framework for supervised learning across VAMP-seq datasets that excludes standard curves. To study the effects of including dataset-specific standard curves downstream of the abundance network during network training (Fig. 2), we also used the model framework shown here to train the abundance network directly against several VAMP-seq datasets in a standard curve-independent manner. The remaining framework modules are identical to the modules of the standard curve-based framework (Fig. 2), and the training procedures are similar.

**Figure S4.**
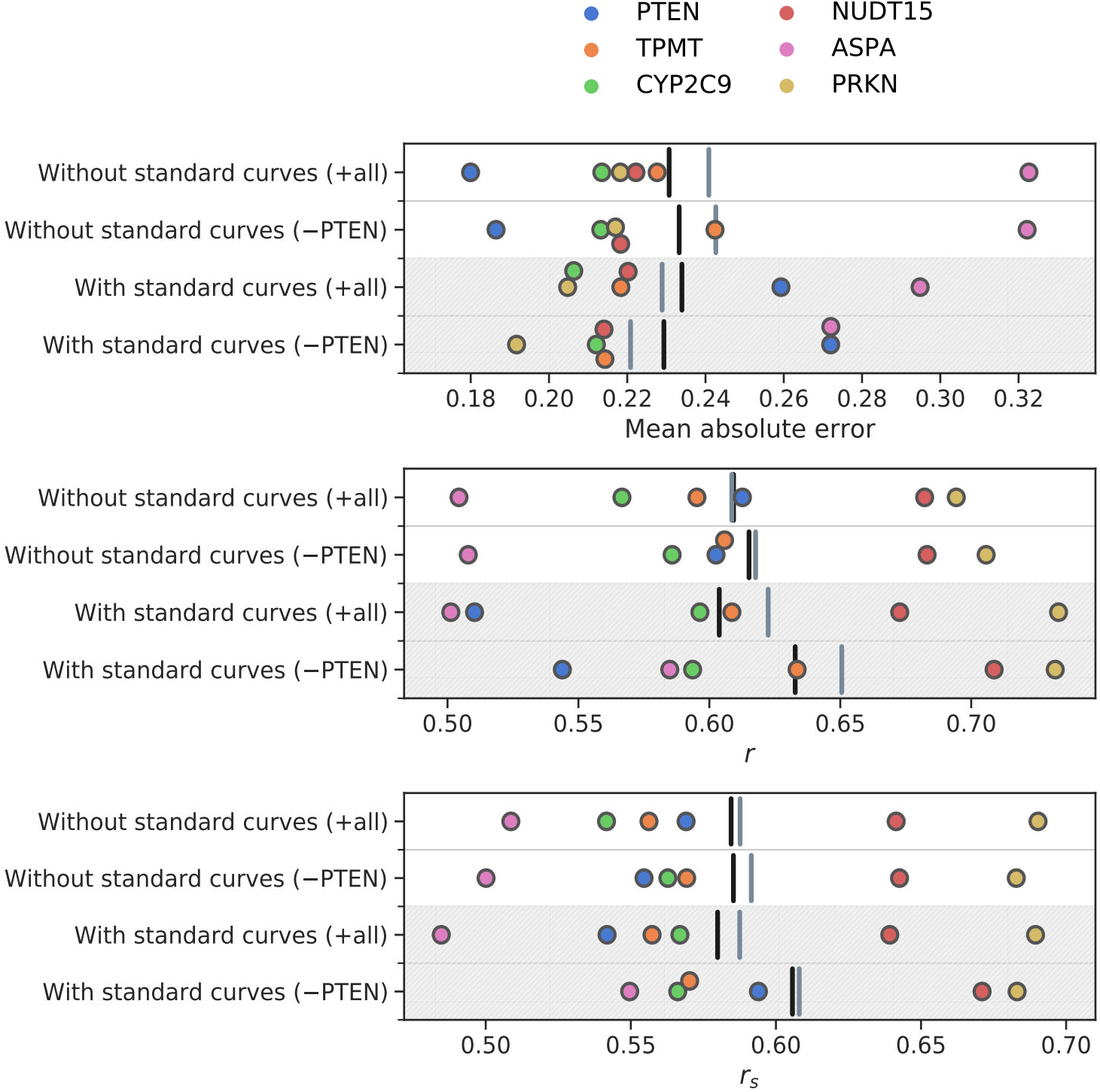
Comparison of performance for models trained without or with standard curves downstream of the abundance network. Leave-one-protein-out cross-validation models were trained without standard curves or with standard curves and either including (+all) or excluding (−PTEN) PTEN VAMP-seq data in the training data. The individual data points indicate the model performance (measured as either mean absolute error, *r* or *r_s_* between predicted and experimental VAMP-seq scores) for a single validation protein for every model type. The black vertical lines indicate the average error or correlation coefficient across all six validation proteins, and the grey vertical lines mark the average error or correlation coefficient across validation proteins excluding PTEN. The models that are here referred to as ‘Without standard curves (−PTEN)’ and ‘With standard curves (−PTEN)’ are called ‘Abundance model (−std. curves)’ and ‘Abundance model (+std. curves)’, respectively, in Fig. 3. Plots that show the same data as here are shown for the individual proteins in Fig. S5.

**Figure S5.**
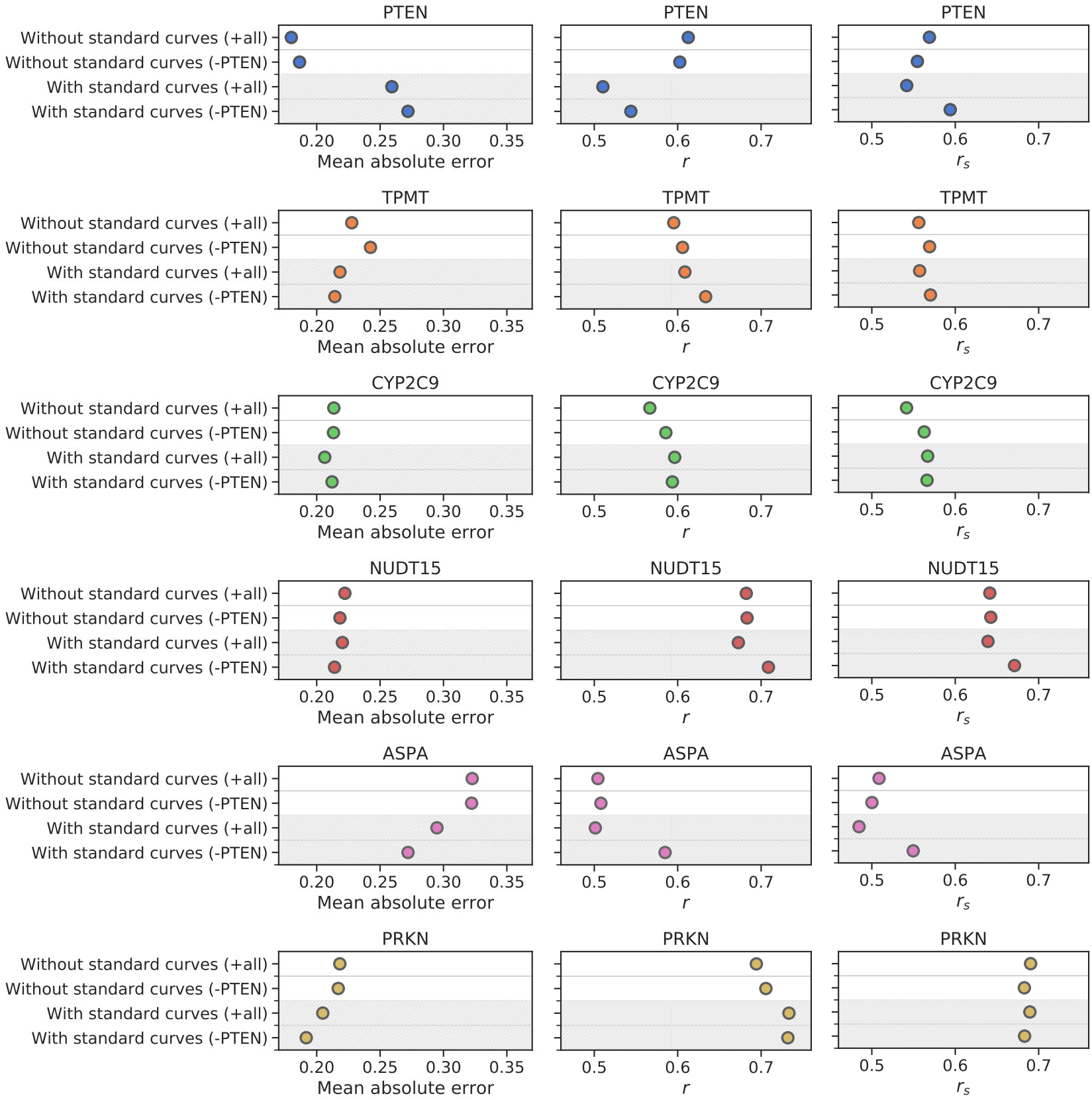
Comparison of performance for models trained without and with standard curves downstream the abundance network. Leave-one-protein-out cross-validation models were trained without standard curves or with standard curves and either including (+all) or excluding (−PTEN) PTEN VAMP-seq data in the training data. The individual data points indicate the model performance (measured as either mean absolute error, *r* or *r_s_* between predicted and experimental VAMP-seq scores) for a single validation protein for every model type. The models that are here referred to as ‘Without standard curves (−PTEN)’ and ‘With standard curves (−PTEN)’ are called ‘Abundance model (−std. curves)’ and ‘Abundance model (+std. curves)’, respectively, in Fig. 3. Averages across model types are shown in Fig. S4

**Figure S6.**
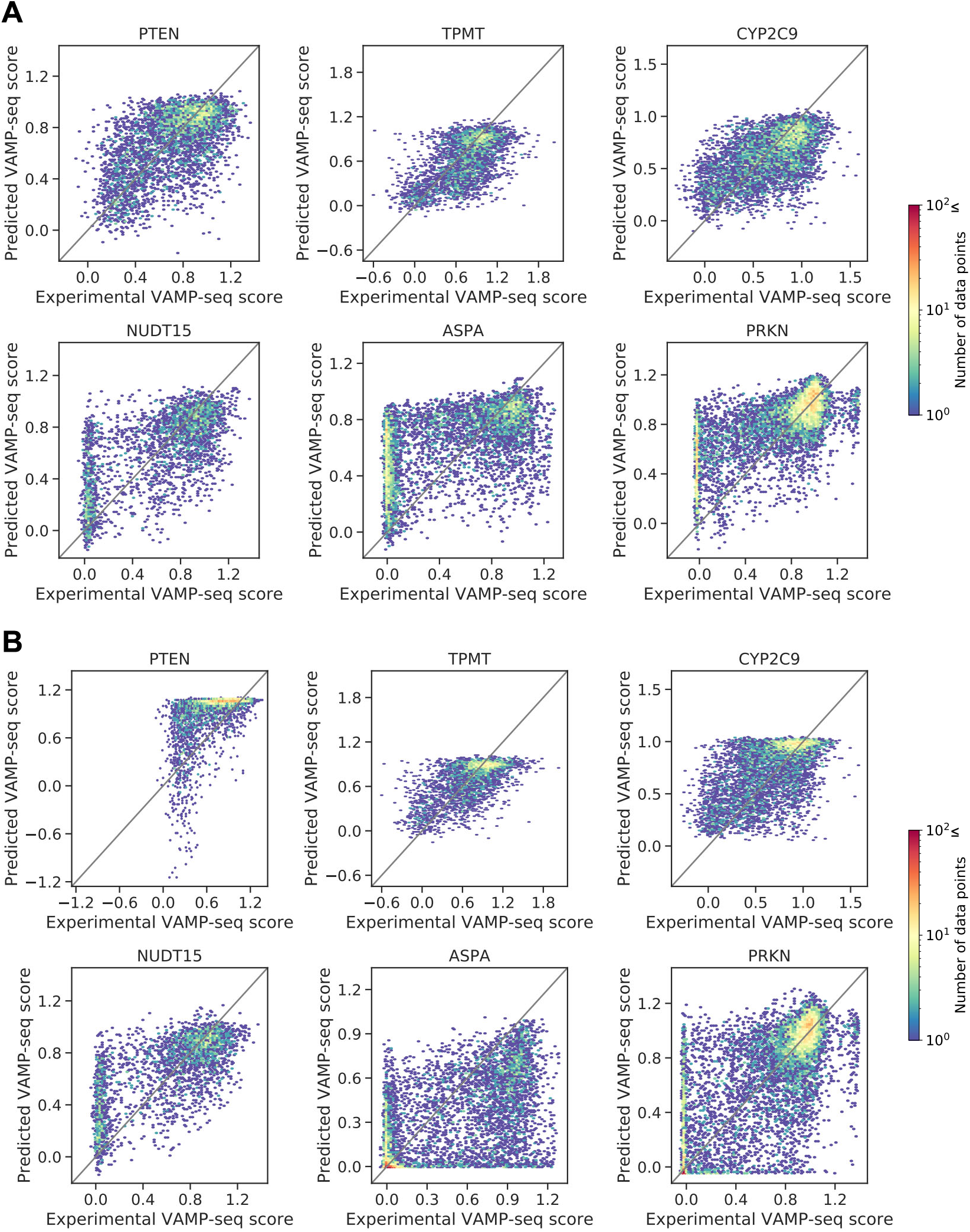
Scatter plots showing predicted VAMP-seq scores as function of experimental VAMP-seq scores for models trained (A) without and (B) with standard curves downstream of the abundance network. Individual scatter plots show prediction results per protein, and abundance score predictions for a given protein always come from a model that was trained without seeing data for that protein. All predictions shown here were moreover generated with models that were trained leaving PTEN VAMP-seq scores out of the training data. Effects of introducing standard curves to the learning framework in particular show up in the PTEN, ASPA and PRKN variant score predictions. The PTEN standard curve maps a wide range of low-throughput scores to VAMP-seq scores around 1, and as a result, many variants have predicted VAMP-seq scores close to 1 when the PTEN standard curve is used. The ASPA standard curve assigns relatively small VAMP-seq scores to variants for which the low-throughput score is WT-like or nearly WT-like. The consequence of this seems to be that VAMP-seq score predictions for many ASPA variants are too small when the ASPA standard curve is applied.

**Figure S7.**
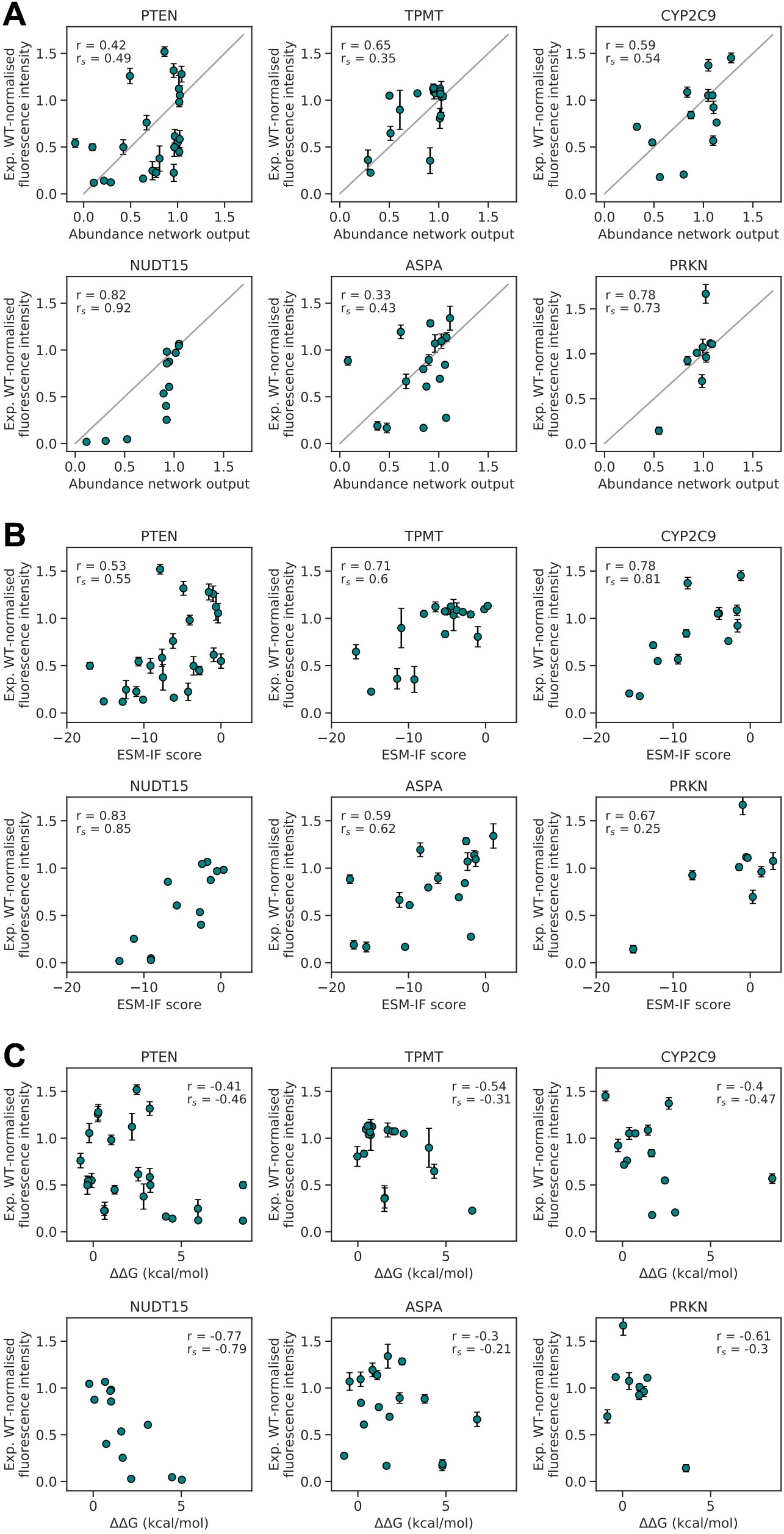
Abundance network and baseline model predictions of WT-normalised GFP:mCherry ratios measured in low throughput. (A) Predictions of low-throughput scores from abundance network (+std. curves, −PTEN) against experimental low-throughput scores. All network predictions come from leave-one-protein-out cross-validation, and neither low- nor high-throughput abundance scores for the validation protein were seen during training. (B) Correlations between low-throughput data and ESM-IF scores. (C) Correlations between low-throughput data and Rosetta ΔΔG values.

**Figure S8.**
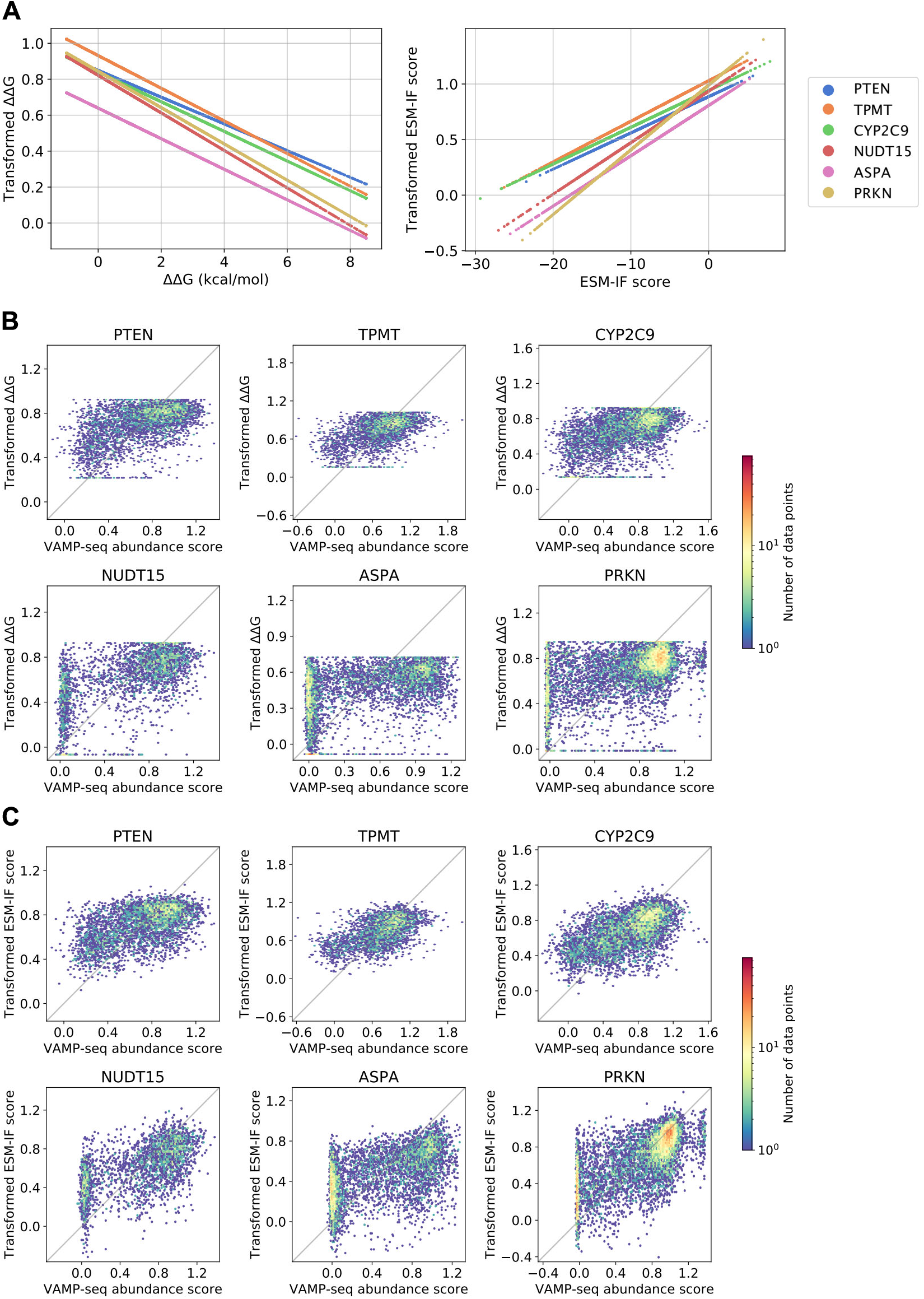
Mapping ΔΔG values and ESM-IF scores to the VAMP-seq score scale. (A) We fit a linear regression model between VAMP-seq scores and either ΔΔG values (left) or ESM-IF scores (right) for each protein in our dataset. We then used the linear models to transform ΔΔG values and ESM-IF scores, thereby obtaining transformed scores falling in the same range as the VAMP-seq abundance scores, but with the same distribition as prior to the transformation. The linear transformations used are here shown for each protein and the two different baseline models. (B) Scatter plots showing experimental VAMP-seq scores vs. ΔΔG values transformed via the linear models. (C) Scatter plots showing experimental VAMP-seq scores vs. ESM-IF scores transformed via the linear models. (B) and (C) thus show the two baseline models as zero-shot predictors of VAMP-seq data.

**Figure S9.**
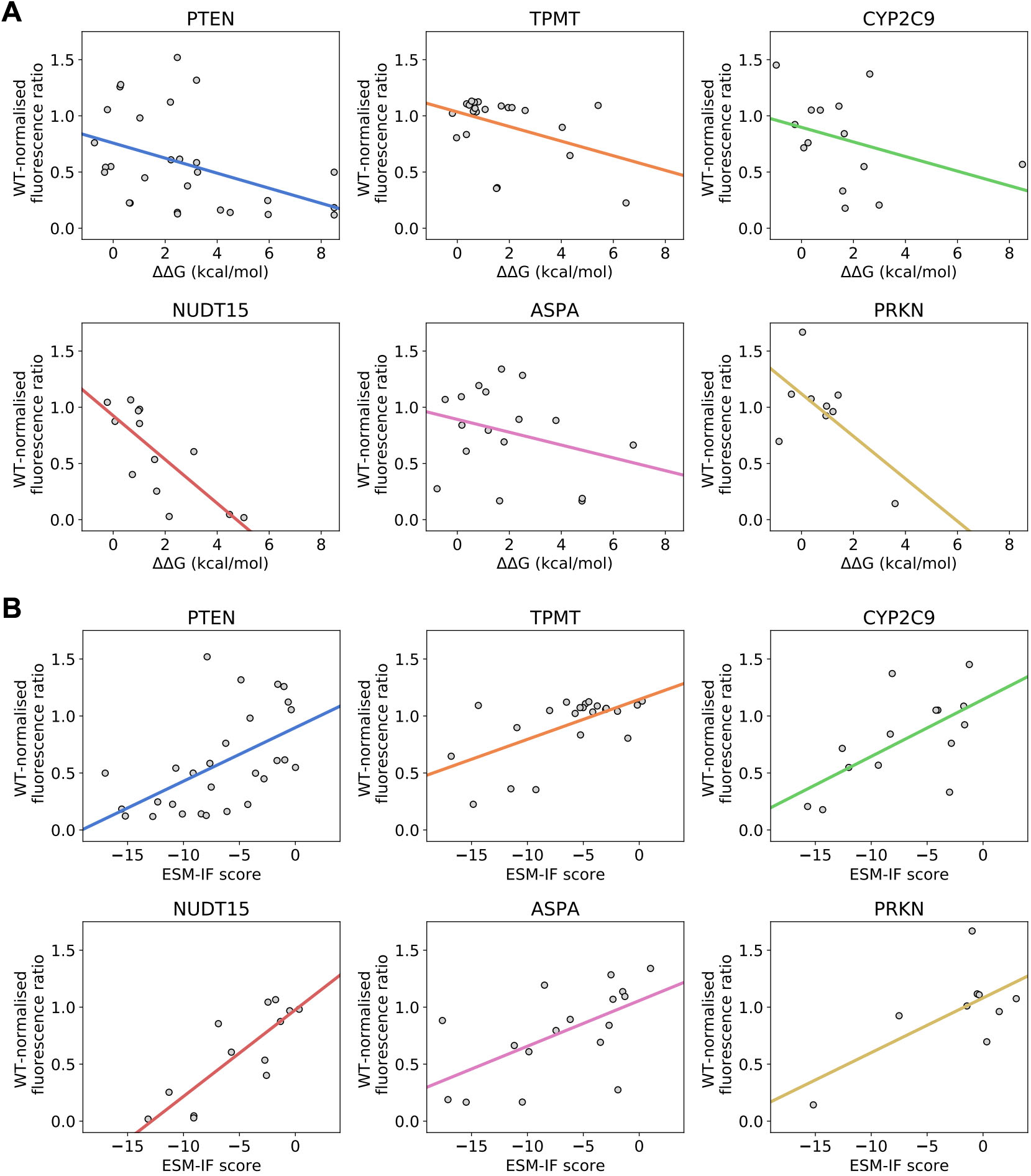
Best linear fits between (A) ΔΔG values or (B) ESM-IF scores and low-throughput measurements of WT-normalised GFP:mCherry fluorescence intensity ratios. The fits were obtained using all available low-throughput data for each protein and used to map ΔΔG values and ESM-IF scores to the low-throughput score space on a per protein-basis.

**Figure S10.**
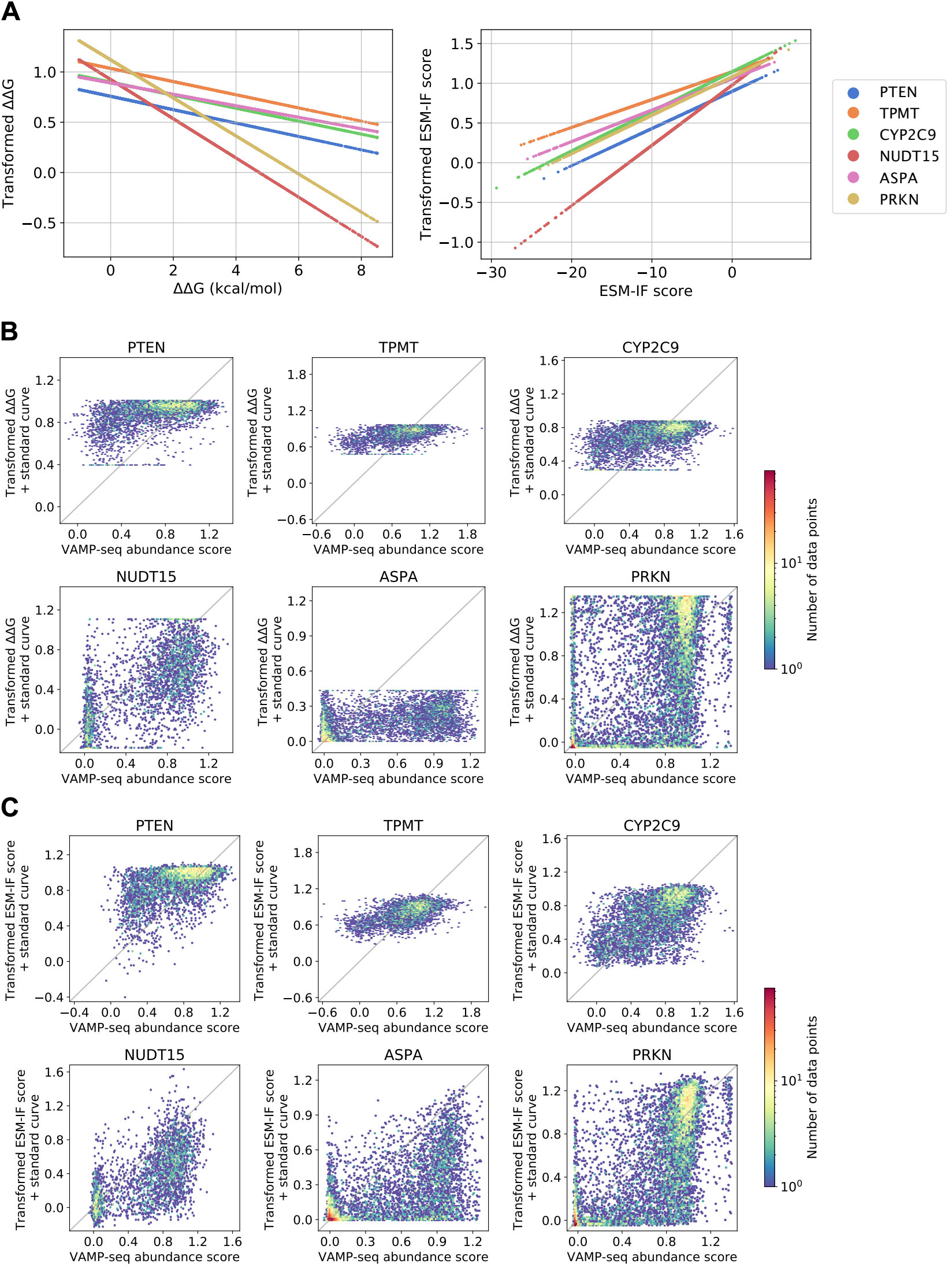
Mapping ΔΔG values and ESM-IF scores to the low-throughput score space followed by transformation with standard curves. (A) We fit a linear regression model between experimental low-throughput scores (Fig. S9) and either ΔΔG values (left) or ESM-IF scores (right) for each protein in our dataset. We then used the linear models to transform ΔΔG values and ESM-IF scores to the low-throughput score space. The linear transformations used are here shown for each protein and the two different baseline models. (B) Scatter plots showing experimental VAMP-seq scores vs. ΔΔG values transformed via the linear models in A and then mapped to the VAMP-seq score space through the standard curves. (C) Scatter plots showing experimental VAMP-seq scores vs. ESM-IF scores transformed via the linear models in A followed by mapping via the standard curves.

**Figure S11.**
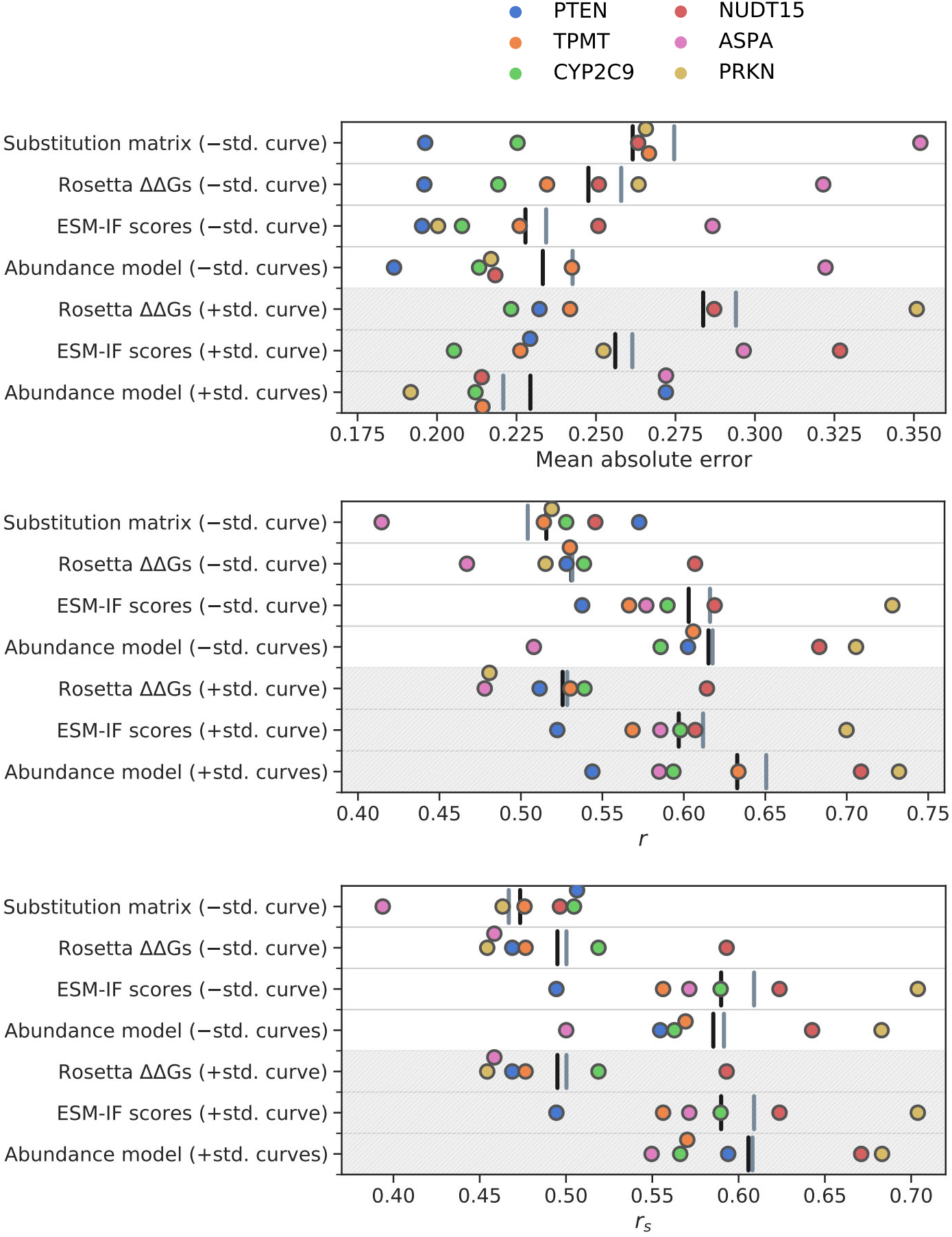
Benchmarking performance of supervised models against all baselines. Comparison of performance of supervised models trained without or with standard curves downstream of the abundance network to the performance of three baseline models. Supervised models were trained without standard curves or with standard curves and excluding PTEN VAMP-seq data from the training data. Baseline model predictions were either mapped linearly to the VAMP-seq score space prior to evaluation (−std. curve) or linearly mapped to the low-throughput score space and then passed through standard curves (+std. curve) before comparison to the experimental data. The individual data points indicate the supervised or baseline model performance (measured as either mean absolute error, *r* or *r_s_* between predicted and experimental VAMP-seq scores) for a single validation protein for every model type. The black vertical lines indicate the average error or correlation coefficient across all six validation proteins, and the grey vertical lines mark the average error or correlation coefficient across validation proteins excluding PTEN. Plots that show the same data as here are shown for the individual proteins in Fig. S12. Areas marked in grey indicate that the predictions have seen the standard curves.

**Figure S12.**
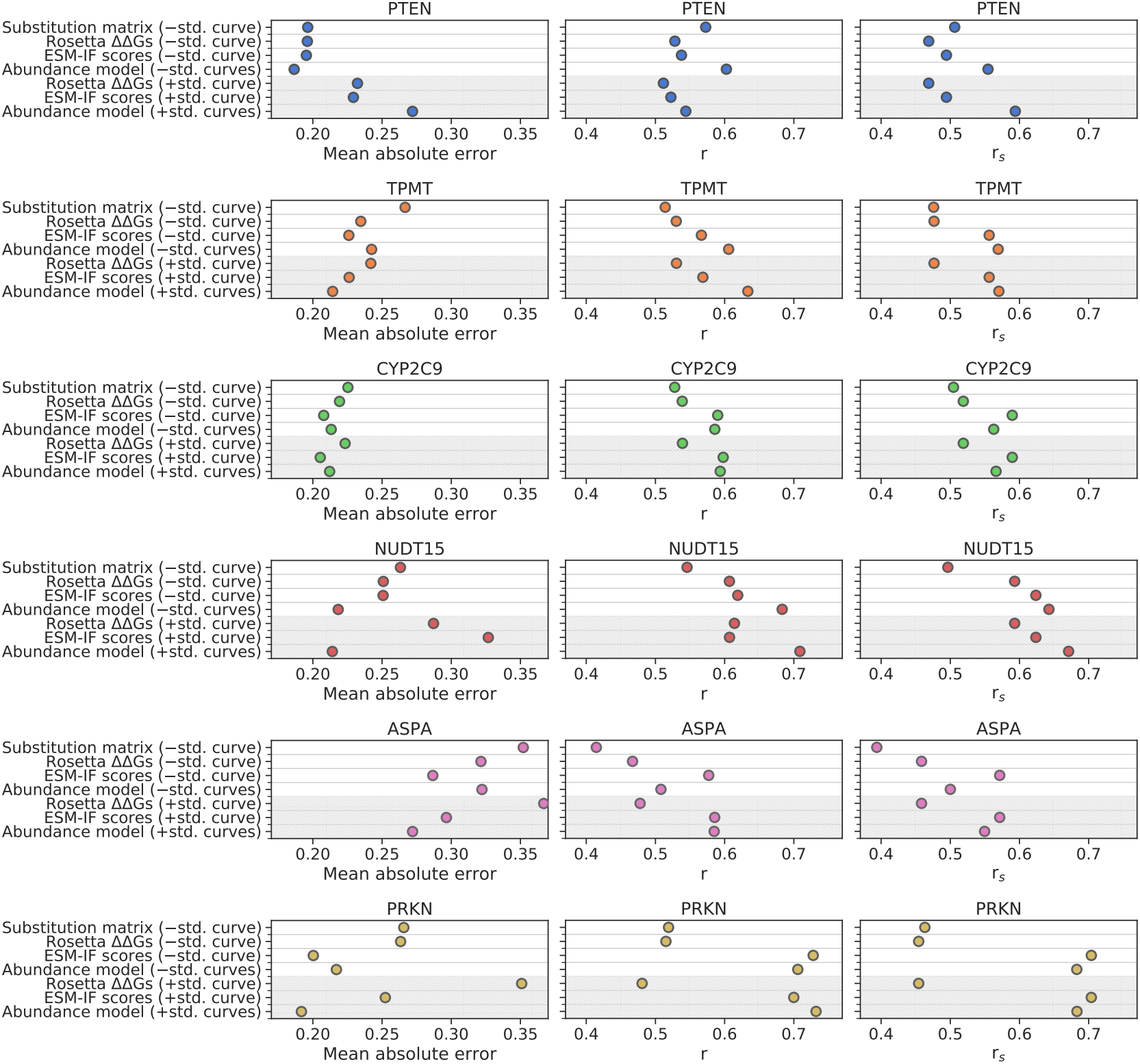
Benchmarking performance of supervised models against all baselines. Comparison of performance of supervised models trained without or with standard curves downstream of the abundance network to the performance of three baseline models. Supervised models were trained without standard curves or with standard curves and excluding PTEN VAMP-seq data from the training data. Baseline model predictions were either mapped linearly to the VAMP-seq score space prior to evaluation (−std. curve) or linearly mapped to the low-throughput score space and then passed through standard curves (+std. curve) before comparison to the experimental data. The individual data points indicate the supervised or baseline model performance (measured as either mean absolute error, *r* or *r_s_* between predicted and experimental VAMP-seq scores) for a single validation protein for every model type. The black vertical lines indicate the average error or correlation coefficient across all six validation proteins, and the grey vertical lines mark the average error or correlation coefficient across validation proteins excluding PTEN. Plots that show the averages across proteins for individual models are shown in Fig. S11. Areas marked in grey indicate that the predictions have seen the standard curves.

**Figure S13.**
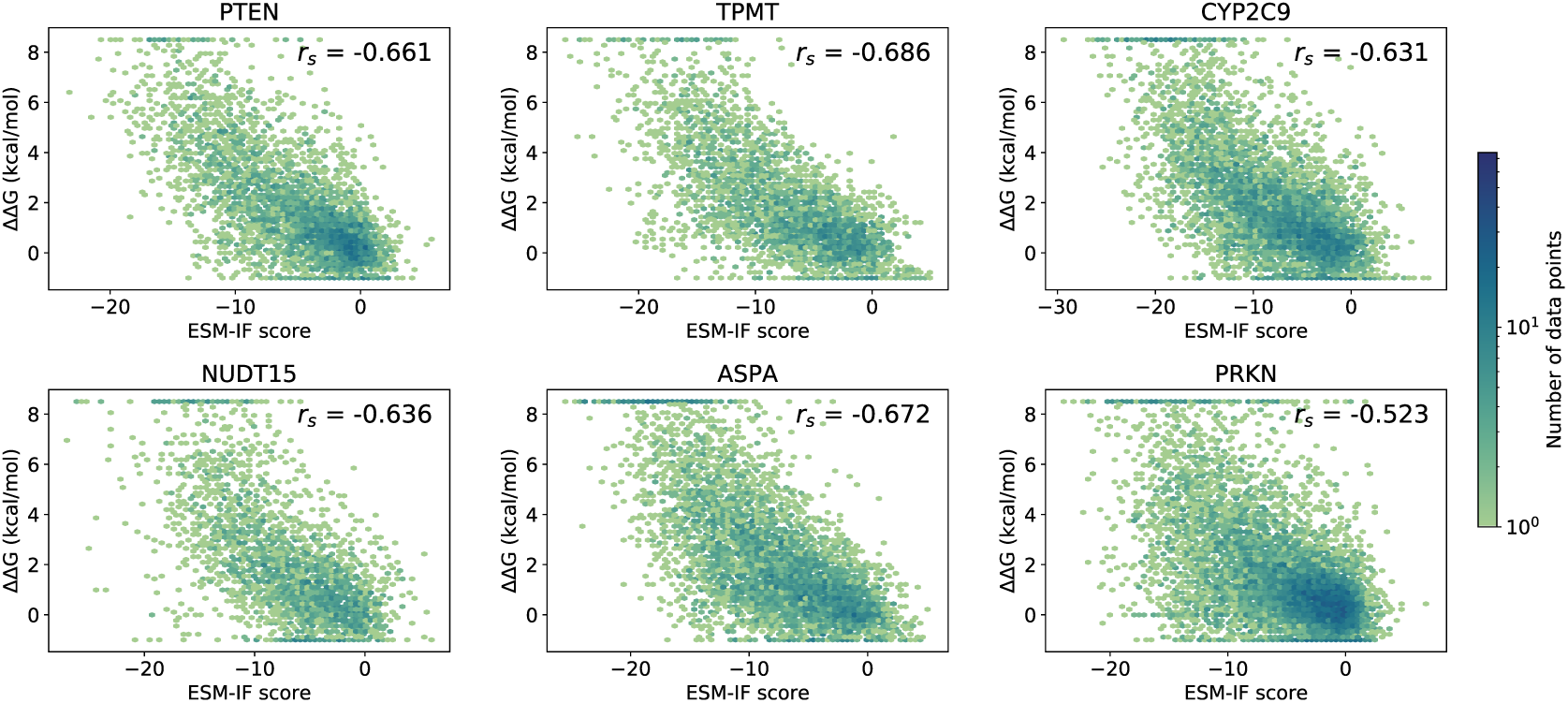
Single residue substitution variant ESM-IF scores and ΔΔG values calculated with Rosetta correlate for individual proteins. The data point density is shown with an upper limit of 75 on the colour scale.

**Figure S14.**
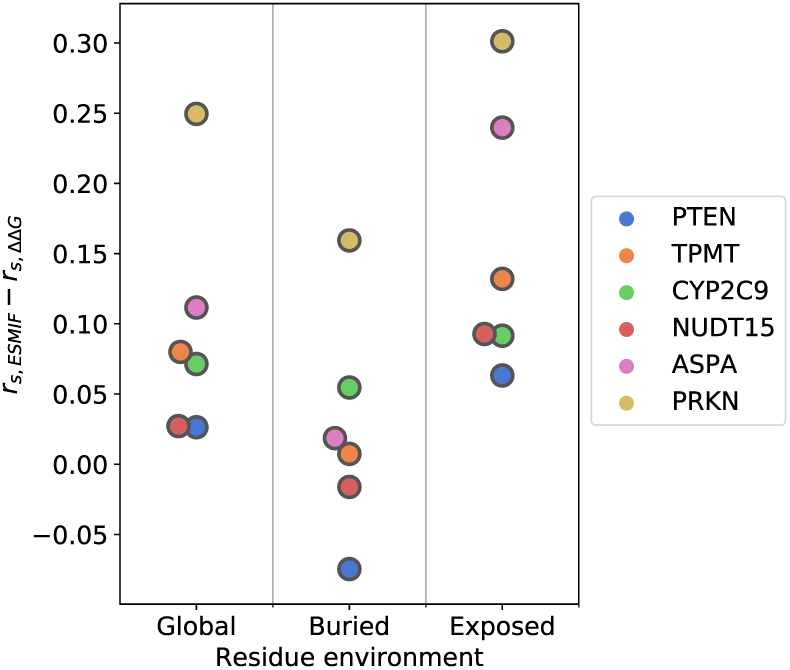
The difference in *r_s_* between VAMP-seq scores and either ESM-IF scores (*r_s,_*_ESM-IF_) or ΔΔG values (*r_s,_*_ΔΔG_) calculated with Rosetta depend on the residue environment. We classified residues as buried or solvent-exposed and calculated *r_s_* between VAMP-seq scores and ESM-IF scores or ΔΔG values for variants of residues in each structure category, and we here show the difference between the two rank correlation coefficients per residue environment. We also show the rank correlation coefficient difference when all residues are considered (Global).

**Figure S15.**
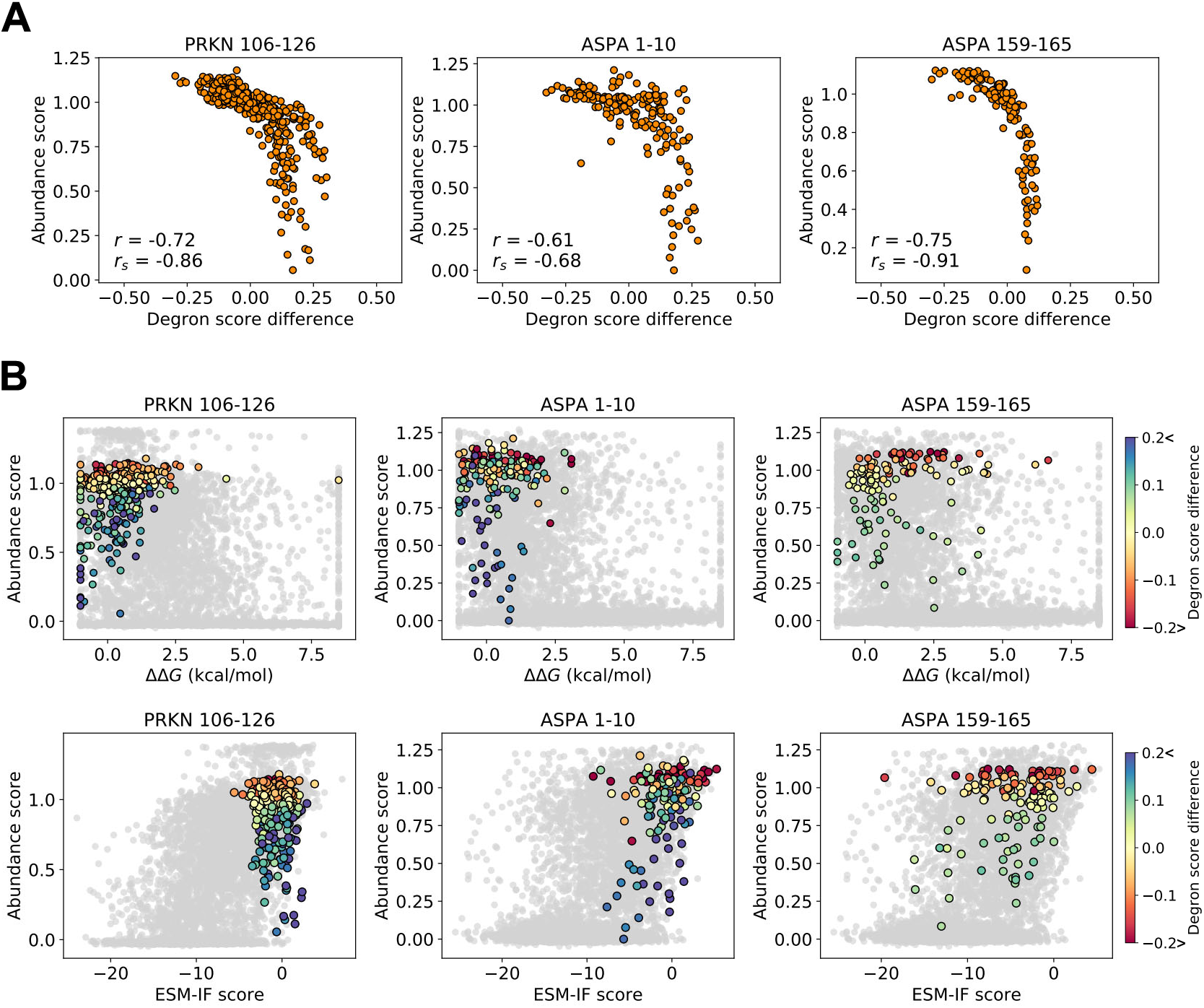
Quality control degron scores correlate with abundance scores and explain some baseline model prediction outliers. (A) We predicted quality control degron scores with QCDpred (***Johansson et al., 2023b***) for wild-type and variant protein sequences and show here how VAMP-seq scores correlate with the differences between variant and wild-type QCDpred scores for all possible single substitution variants in selected regions of PRKN and ASPA. The side chains of residues in the selected regions are all relatively solvent-exposed. As previously observed, the abundance scores of PRKN variants in residues 106–126, which belong to a disordered loop region in the protein, show high correlation with QCDpred score differences (***Clausen et al., 2024***). Abundance scores of variants of the ASPA N-terminus (residues 1–10) and an ASPA loop region (residues 159-165) are similarly highly correlated with predicted degron propensity differences between variant and wild-type residues. (B) Scatter plots of abundance scores and either ΔΔG values (top) or ESM-IF scores (bottom) for PRKN and ASPA variants. Data for all variants in the VAMP-seq datasets are shown in light gray, and variants of PRKN residues 106–126 and ASPA residues 1–10 or 159–165 are highlighted and coloured according to the predicted difference between variant and wild-type degron propensities. QCDpred score differences might explain why some variants reduce cellular abundance, even though they are not predicted to thermodynamically destabilise the proteins (ΔΔG ≈ 0 kcal/mol) or to reduce the variant score predicted by ESM-IF. Moreover, variants with wild-type-like abundance scores, but predicted structural destabilisation, might be explained by the fact that the relevant substitutions decrease the local degron propensity.

**Figure S16.**
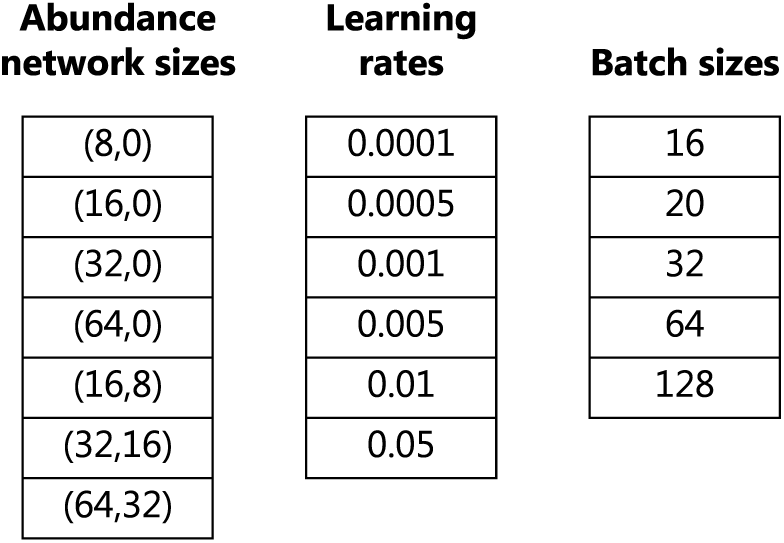
Overview of hyperparameters optimised for supervised models. We scanned abundance network sizes, learning rates and batch sizes to optimise the performance of our trained models. The hyperparameter search space was defined by the values listed for each hyperparameter here. Based on these lists, we randomly generated 50 combinations of the three hyperparameters and trained models with these 50 random hyperparameter combinations. We always scanned the same 50 randomly sampled combinations of the three hyperparameters and report the best combinations for each type of trained model in Table 1.

**Figure S17.**
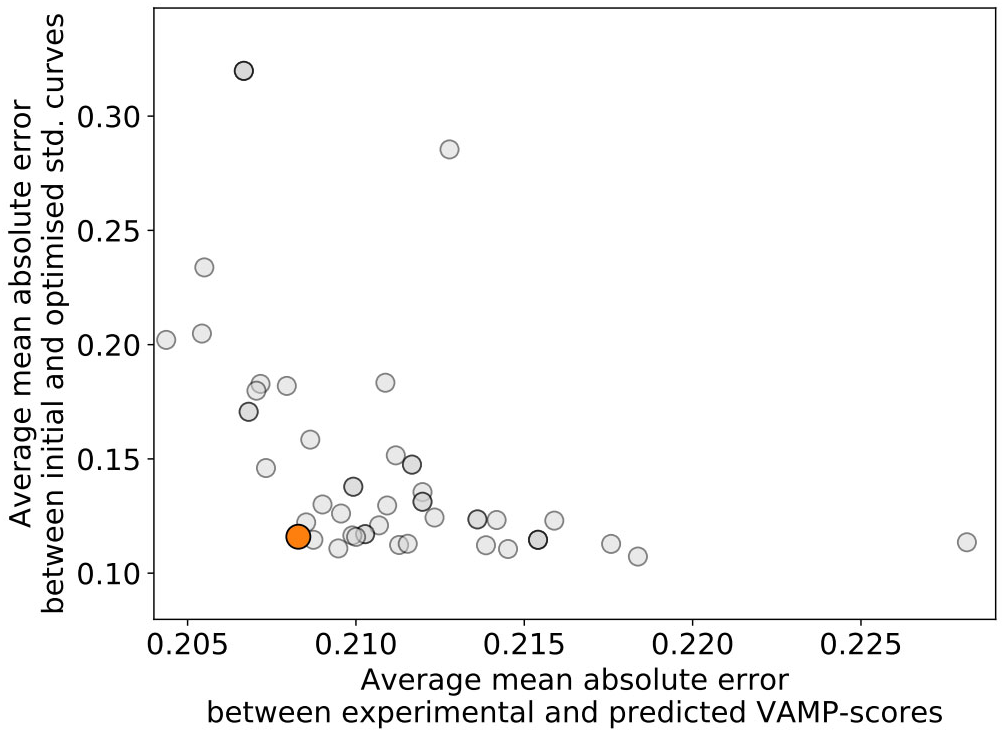
Results of hyperparameter scan for models trained with standard curve fine-tuning. We experimented with fine-tuning of the standard curves during abundance model training and trained models using leave-one-protein-out cross-validation for 50 combinations of selected hyperparameters (Fig. S16). Each data point in the scatter plot shows the results for one of the 50 hyperparameter combinations, more specifically the mean absolute error between predicted and experimental VAMP-seq scores averaged across validation proteins and the mean absolute error between initial and fine-tuned standard curves averaged across proteins for that single hyperparameter combination. Above, we report results for the hyperparameter set highlighted in orange (see details in Table 1), since this parameter set minimises the average validation error while simultaneously minimising the average shift in standard curves away from the initial fits.

**Figure S18.**
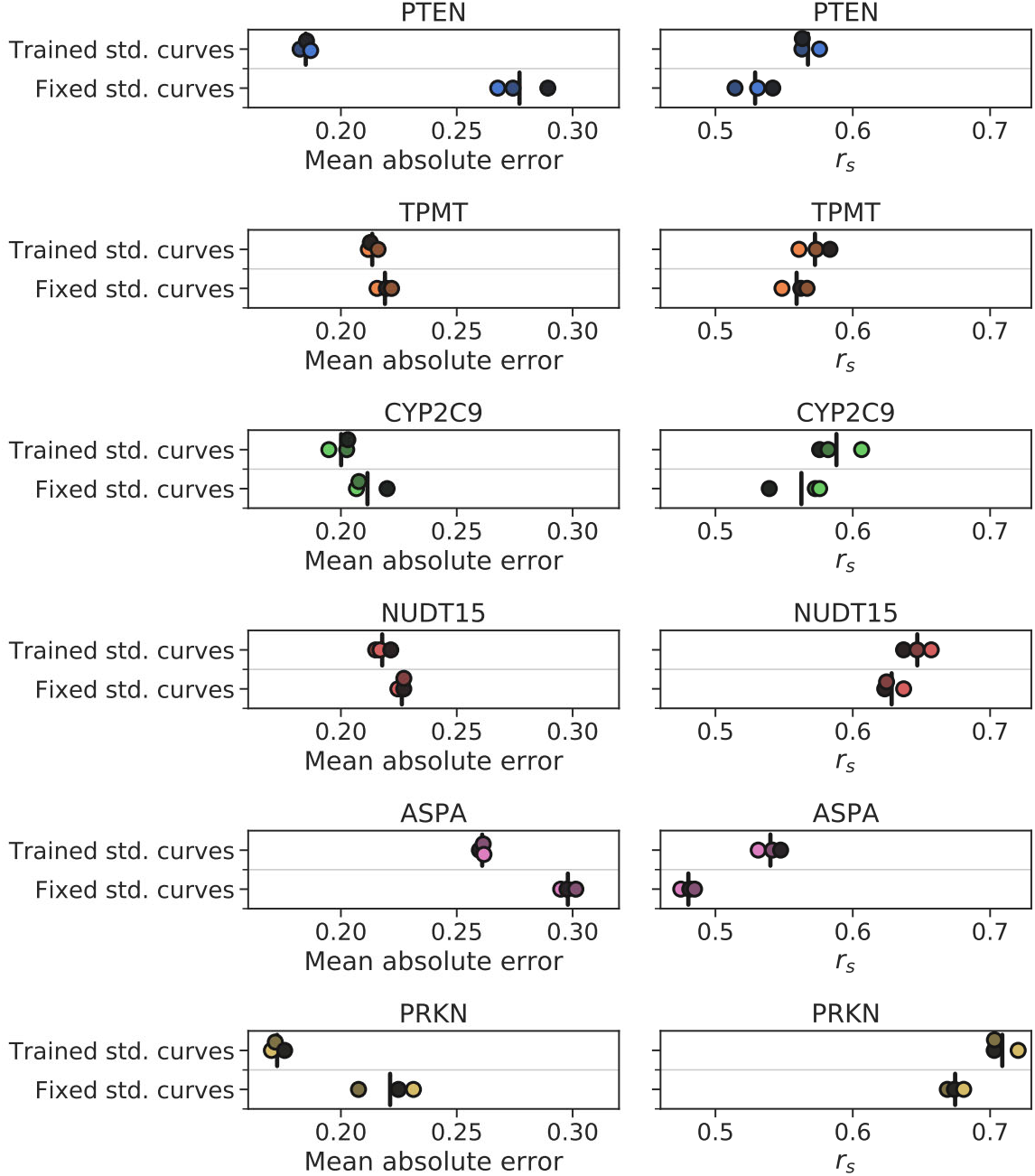
Fine-tuning of standard curves during abundance model training improves model performance. The mean absolute error and *r_s_* between experimental and predicted VAMP-seq scores are shown for abundance models trained with trainable standard curves and abundance models trained with fixed standard curves. Training was performed using leave-one-protein-out cross-validation, and all available VAMP-seq datasets were included in the cross-validation. Models were trained in ensembles of three, and individual data points show results for individual models in the ensembles. The performance of individual models were evaluated on 80% of variant effect scores for the validation protein, since 20% of scores were used to adjust the validation protein standard curve during training with trainable standard curves. Vertical black lines indicate the average across the three individual evaluations.

**Figure S19.**
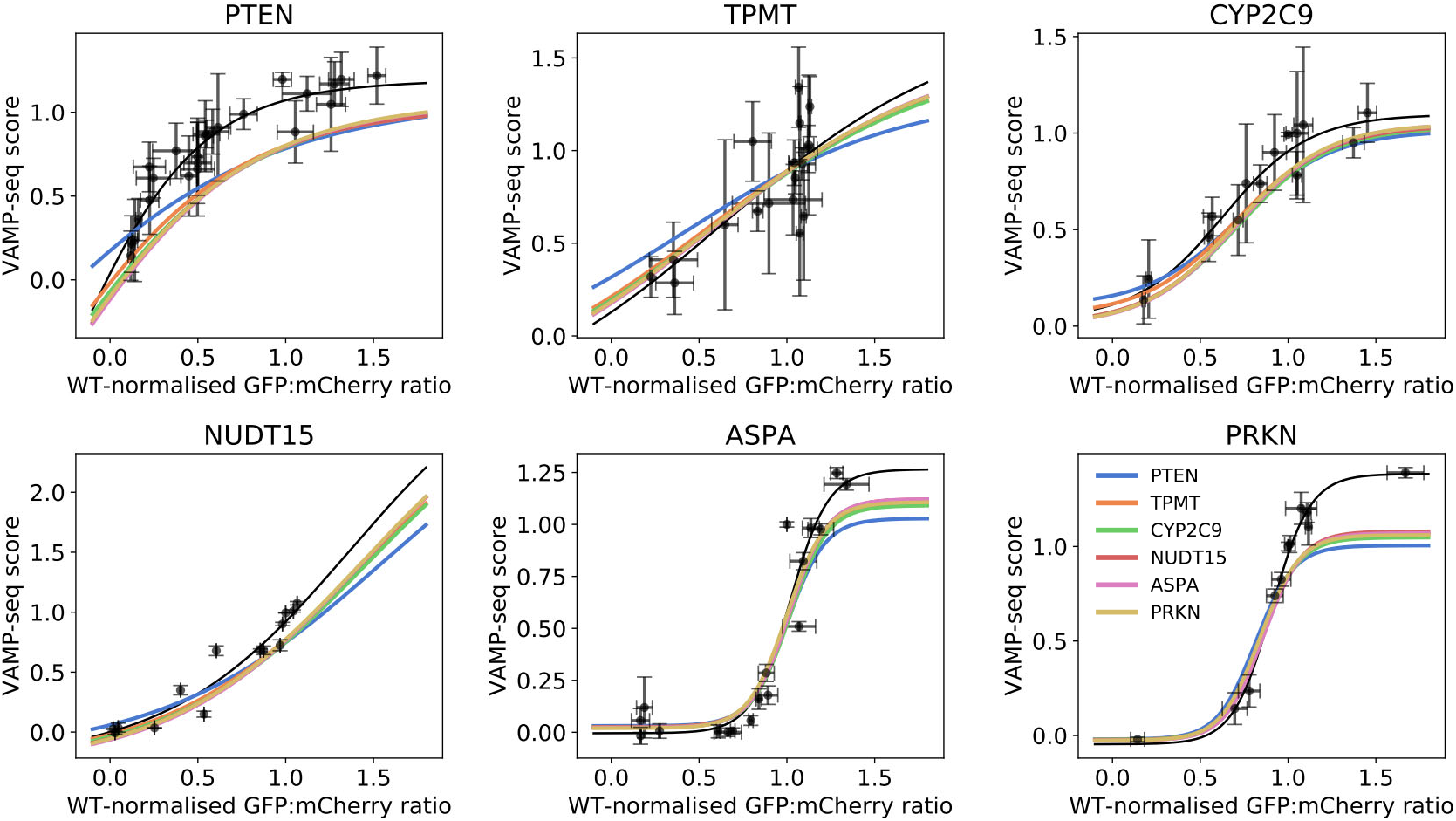
Standard curves before and after fine-tuning during abundance model training. The initial standard curves, which correspond to the standard curves in Fig. 1, are shown in black together with the experimental data that was used to fit the curves. For each protein, the standard curves that were obtained after abundance model training that included standard curve fine-tuning are shown in various colours. The six coloured curves specifically correspond to the six curves that were obtained when each of the six proteins was used as validation protein in the leave-one-protein-out cross-validation procedure. In the PTEN plot, the blue standard curve is for example the standard curve resulting from abundance model training and standard curve fine-tuning with PTEN as validation protein, and the orange curve in the same plot is the standard curve obtained with TPMT as validation protein. We observe that the six curves for each protein are relatively similar, in particular in areas with experimental information (black data points). The coloured standard curves shown here are averaged across curves from the individual base models in our three model-ensembles.

